# *O*-GlcNAcylation is essential for rapid *Pomc* expression and cell proliferation in corticotropic tumor cells

**DOI:** 10.1101/2021.08.13.455965

**Authors:** Logan J Massman, Michael Pereckas, Nathan T Zwagerman, Stephanie Olivier-Van Stichelen

## Abstract

Pituitary adenomas have a staggering 16.7% lifetime prevalence and can be devastating in many patients due to profound endocrine and neurologic dysfunction. To date, no clear genomic or epigenomic markers correlates with their onset or severity. Herein, we investigate the impact of the *O*-GlcNAc post-translational modification in their etiology. Found in over 5000 human proteins to date, *O*-GlcNAcylation dynamically regulates proteins in critical signaling pathways, and its deregulation is involved in cancers progression and endocrine diseases such as diabetes.

In this study, we demonstrate that *O*-GlcNAcylation enzymes were upregulated, particularly in aggressive ACTH-secreting tumors, suggesting a role for *O*-GlcNAcylation in pituitary adenoma etiology. In addition to the demonstration that *O*-GlcNAcylation was essential for their proliferation, we show that the endocrine function of pituitary adenoma is also dependent on *O*-GlcNAcylation. In corticotropic tumors, hyper-secretion of the proopiomelanocortin (POMC)-derived hormone ACTH leads to Cushing’s disease, materialized by severe endocrine disruption and increased mortality. We demonstrate that *Pomc* mRNA is stabilized in an *O*-GlcNAc-dependent manner in response to corticotropic-stimulating hormone (CRH). By impacting *Pomc* mRNA splicing and stability, *O*-GlcNAcylation contributes to this new mechanism of fast hormonal response in corticotropes. Thus, this study stresses the essential role of *O*-GlcNAcylation in ACTH-secreting adenomas’ pathophysiology, including cellular proliferation and hypersecretion.

## INTRODUCTION

Approximately 16.7% of people develop a pituitary adenoma in their lifetime (2). Consequently, around 10,000 pituitary adenomas are diagnosed yearly in the US alone, accounting for 15% of central nervous system tumors (2–4). The pathophysiology of pituitary adenomas is diverse and are defined clinicopathologically as one of the following subtypes: adrenocorticotropic hormone (ACTH)-, growth hormone (GH)-, thyroid-stimulating hormone (TSH)-, and gonadotropin (luteinizing hormone (LH)/ follicle-stimulating hormone (FSH))- and prolactin (PRL)-secreting, and non-secreting. Secreting adenomas present subtype-specific clinical symptoms. For example, GH-secreting tumors may induce acromegaly, leading to physical impairment, body pain, a significantly reduced quality of life, and increased mortality (3,4). On the other hand, ACTH-secreting adenomas cause Cushing’s disease, associated with increased mortality (5). Finally, PRL-secreting adenomas are associated with amenorrhea, galactorrhea, sexual dysfunction, and infertility (6). The other subtypes of pituitary adenomas, including non-secreting subtypes, may lead to hypopituitarism by compressing surrounding tissue, decreasing patient quality of life, and increasing the risk of cardiovascular events and premature death (4,7). In addition, pituitary adenomas may also compress the optic chiasm, leading to visual defects (10) or invade neighboring dural structures in more than 50% of patients (11,12). Therefore, even in the absence of hormonal disruption, these neurologic dysfunctions grandly impact patients’ quality of life and requires medical care.

Medical treatment for functional pituitary tumors is an ever-growing field. Current medical management of acromegaly includes octreotide acetate, which often requires lifelong treatment and confers a significant financial burden (13). Likewise, medical management for Cushing’s disease often focuses on blocking the downstream effects of cortisol rather than treating the tumor itself (14). The lack of mechanistic insight thus limits the treatment protocols (15), with surgery being the main treatment for most pituitary adenomas. However, resection is particularly challenging for small microadenomas with dramatic clinical sequelae, such as Cushing-associated adenomas, necessitating arduous resection, often associated with higher morbidity rate (12). Therefore, increasing our molecular understanding of pituitary adenoma pathophysiology is essential to find better targeted treatments and improve clinical outcomes.

Despite their prevalence and significant associated morbidities, the tumorigenesis of pituitary adenomas remains poorly understood. Although a small subset of pituitary adenoma cases is familial (most often associated with *MEN1* mutations), the vast majority are sporadic with no clear common etiology (8,17,18). In addition, the heterogeneity of pituitary adenomas, emerging from diverse endocrine cell types, is highly challenging and requires targeted studies to understand the specific molecular signaling pathways dysregulated in each tumor subtype. In this article, we focused on ACTH-secreting pituitary adenomas due to their potential for profound endocrine disruption and lack of medical therapies targeted at their underlying pathophysiology.

While pituitary adenomas have been studied for decades, much remains to be learned about their tumorigenesis, growth, and function. Whole-exome sequencing of pituitary adenomas demonstrated a low number of somatic mutations not involved in their tumorigenic potential (14). Next-generation sequencing of the mitochondrial genome in pituitary adenomas revealed many novel variants in these tumors, again not correlating with clinicopathological features (15). While DNA methylation and histone acetylation markers have been linked to invasiveness, the functional role of epigenetic changes in the endocrine and proliferative components of pituitary adenomas has not been studied further (16). Taken together, our understanding of tumorigenesis and hypersecretory function of pituitary adenomas is highly limited when viewed through a genetic lens, and more diverse non-genetic mechanisms must therefore be explored to pave the way to developing novel treatment protocols.

With this goal in mind, we questioned the importance of the *O*-GlcNAc post-translational modification in pituitary tumor development. In short, this modification consists of the reversible addition of a single sugar residue (N-acetylglucosamine) onto serine or threonine of nuclear, cytoplasmic, and mitochondrial proteins (Figure 1). Two essential enzymes regulate this cycling – *O*-GlcNAc Transferase (OGT) and *O*-GlcNAcase (OGA) – which reversibly add or remove *O*-GlcNAc residues, respectively. With more than 5000 *O*-GlcNAcylated human proteins described to date and counting (21), these post-translationally modified proteins are involved in essential signaling processes such as transcription/translation, cell cycle, metabolism, cell survival, and protein degradation (21). Highly dependent on extracellular glucose concentrations, this nutrient-sensing modification adapts cell signaling in response to changes in nutrient availability (22). Finally, *O*-GlcNAcylation is dysregulated in many pathological conditions such as neurodegenerative and cardiovascular diseases and cancers (23–25).

**Figure 1:**
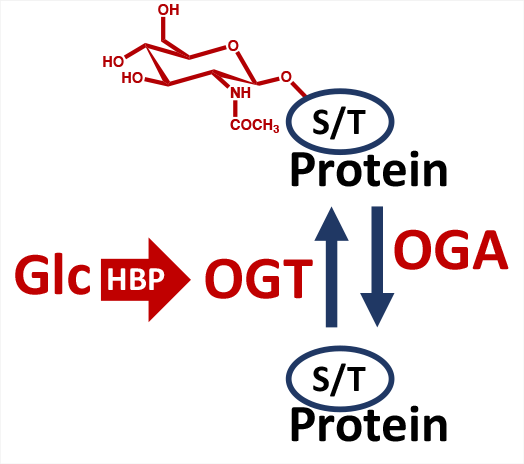
O-GlcNAcylation of proteins. 2-3% of glucose (Glc) entering cells is shuttled through the Hexosamine Biosynthesis Pathway (HBP) to produce UDP-GlcNAc, the nucleotide donor for O-GlcNAcylation. O-GlcNAc transferase (OGT) adds the GlcNAc residue to serine or threonine (S/T) residues. O-GlcNAcase (OGA) removes the sugar residue.

Increased overall *O*-GlcNAcylation levels are consistently observed in cancer cells, including breast, lung, and colorectal (26,27). Further, *O*-GlcNAcylation competes with phosphorylation sites on oncogenes and tumor suppressors to increase proliferation and promote invasion, making it an essential mediator in tumor formation and progression (20–23). Finally, ACTH-secreting pituitary adenomas increase glucose transporter’s cell surface expression in response to corticotrophin-releasing hormone (CRH)(30), increasing glucose uptake and presumably *O*-GlcNAcylation levels. Despite this, the involvement of *O*-GlcNAc cycling has never been investigated in pituitary tumors. We, therefore, hypothesize that *O-* GlcNAcylation contributes to pituitary tumor development. More precisely, we believe that *O*-GlcNAcylation impacts multiple aspects of these tumors, including their hyper-proliferative and hyper-endocrine abilities.

In this study, we used a combination of normal and tumorous human pituitary tissue and the AtT-20 murine pituitary adenoma cell line to study the role of *O*-GlcNAcylation in pituitary adenoma etiology. We demonstrated that *O*-GlcNAc cycling enzymes *OGT* and *OGA* are significantly upregulated in ACTH-secreting tumors, correlating with the aggressiveness and ACTH secretion levels. Accordingly, *O*-GlcNAcylation was essential for pituitary adenoma cells’ proliferation, a common function for *O*-GlcNAcylation in both development and adult cells. This finding identifies *O*-GlcNAcylation as a potential therapeutic target in pituitary adenomas. As the predominant cause for clinical intervention, we next investigated pituitary adenoma hypersecretory ability. In AtT-20 ACTH-secreting pituitary adenoma-derived cells, the endocrine function is characterized by the expression and secretion of proopiomelanocortin (POMC)-derived hormones upon CRH stimulation. For the first time, we demonstrated that CRH transiently stabilizes the pool of available *Pomc* mRNA to quickly raise *Pomc* levels, independently of the *Pomc* promoter activation consequences of the CRH-downstream signaling pathway. Interestingly, this mechanism exists for many hormones, including TSH, luteinizing hormone (LH), and follicle-stimulating hormone (FSH) mRNA (31–36), and has been suggested for POMC pre-hormone as well (37). Furthermore, *O*-GlcNAcylation is essential for this novel regulatory pathway as it enhances the proper splicing of *Pomc* mRNA and prevents its decay. Finally, we present a list of *O*-GlcNAcylated targets responsive to CRH stimulation and involve in mRNA processing. While those results are preliminary, we are hopeful that the expansion of this research in pituitary adenoma subtypes and more extensive clinical samples will highlight some of these *O*-GlcNAcylated proteins as druggable targets for corticotropic pituitary adenomas treatment.

## MATERIALS AND METHODS

### Normal pituitary gland and pituitary adenoma collection

A terminally ill 74-year-old male amyotrophic lateral sclerosis (ALS) patient was enrolled for post-mortem brain/pituitary collection under an IRB-approved study. To date, ALS ontology is not correlated with the pituitary gland’s molecular alteration or *O*-GlcNAc defect (38), and no abnormality was noted on the pituitary at the autopsy. The pituitary gland was flash-frozen upon collection and used as normal pituitary tissue. Patients with pituitary adenomas were consented before surgery under an IRB-approved protocol. Pituitary adenomas were frozen and stored promptly following the resection. Patients were chosen based on the availability of a sufficient amount of banked tissues post-pathology analysis to create a representative panel of the various pituitary adenoma subtypes. Mammosomatotrophic tumors were characterized by GH+/PRL+ on histology, corticotropic tumors by ACTH+ on histology, and non-secreting tumors by negative hormone staining. Cushing’s disease was defined by signs and symptoms of cortisol excess and elevations of late-night salivary cortisol levels on two separate samples. Furthermore, if imaging was equivocal, inferior petrosal sinus sampling was performed to confirm pituitary etiology and provide guidance for laterality for surgical planning. Acromegaly patients were defined by signs and symptoms of growth hormone excess followed by lab values indicating elevations of IGF-1 levels and then confirmed with a growth hormone suppression test in which a failure to suppress was indicative of acromegaly. Pre-operative clinical diagnostic and pathological data for each subject is presented in Table S1(39). Two non-secreting, 2 mammosomatotrophic (1 with acromegaly at recurrence) and 4 corticotropic tumors (3 Cushing’s disease and 1 hypopituitarism pre-operative clinical diagnosis) were analyzed in this study.

### Cell culture

AtT-20 cells were maintained in Dulbecco’s modified Eagle’s medium (DMEM) containing either 4.5 g/L (Standard, Std) or 1g/L (Low) Glucose, 10% (v/v) fetal bovine serum (FBS), and 1% penicillin/streptomycin at 37°C in a 5% (v/v) CO_2_-enriched humidified atmosphere. Cells were cultivated in Std or Low glucose media for at least 24h before the start of the experiments.

### Drugs and siRNA

Lyophilized corticotropin-releasing hormone (CRH, Sigma) was reconstituted in phosphate-buffered saline (PBS) with 10 mM acetic acid for stability and used at a final concentration of 28 μM. Thiamet G (100 nM final concentration) and 5S-GlcNAc (100 μM final concentration) were dissolved in sterile distilled water. 5,6-dichlorobenzimidazole 1-beta-D-ribofuranoside (DRB) (50 μM final concentration) and OSMI4 (5μM final concentration) were resuspended in DMSO, and corresponding control samples were treated with a similar amount of DMSO. Cells were treated with inhibitors for 16h unless otherwise precise. Mission® esiRNA targeting mouse *Oga* (Mgea5) was purchased from Sigma (#EMU020941). Silencer™ Negative Control siRNA was also purchased from Sigma (#AM4611). One microgram of siRNA was transfected in one well of a 6-well using Lipofectamine RNAiMAX transfection reagent according to the manufacturer protocol (Thermofisher Scientific). Cells were treated with siRNA for 24h.

### RNA extraction, cDNA synthesis, PCR, and qPCR

mRNA was isolated with the PureLink RNA Mini Kit (#12183018A, Thermo Fisher Scientific) following the kit instruction. RNA concentrations were measured using an LVis microplate (BMG Labtech). cDNA was then synthesized with SuperScript™ IV VILO™ Master Mix with ezDNase (#11766050, Thermofisher Scientific) according to the manufacturer’s instructions. The resulting cDNA was diluted sequentially to a 1:100 dilution and used as templates for PCR.

PCR was performed using DreamTaq Green PCR Master Mix (Thermofisher Scientific) and loaded on 2% agarose gels in TAE buffer. qPCR reaction was then performed using PowerSYBR qPCR Master Mix (Thermo Fisher Scientific). Specific primers for each reaction are as follows: *Pom Ex-Ex*-F: GGAGAGAAAGCCGAGTCACAA; Pomc *Ex-Ex*-R: GGGACCCCGTCCTGTCCTATAA; *Pomc Intron*-F: TTCGTTCATTGGAGTGGCCC; Pomc Intron-R: CGTACTTCCGGGGGTTTTCA; A*ctin*-F: AGATCAAGATCATTGCTCCTCCT; Actin-R: ACGCAGCTCAGTAACAGTCC; Og*t*-F: TTCGGGAATCACCCTACTTCA; *Ogt*-R: TACCATCATCCGGGCTCAA; *Oga-*F: CGGTGTCGTGGAAGGGTTTTA; *Oga-*R: GTTGCTCAGCTTCTTCCACTG. qPCR was performed on a QuantStudio 3 instrument (Applied Biosystems) with the recommended settings. The data were collected and processed using DataConnect (Thermo Fisher Scientific), and 2^−ΔΔCq^ were calculated and plotted using Prism 9 (GraphPad Software).

### Protein lysis, SDS-PAGE, and Western Blotting

Samples were lysed for 5 min in RIPA lysis buffer [10mM Tris-HCl, 150mM NaCl, 1% Triton X-100 (v/v), 0.5% NaDOC (w/v), 0.1% SDS (w/v), and protease inhibitors; pH7.5], vortexed and centrifuged 10 minutes at 18,000 g at 4°C. Sample lysates were resolved on 8% Tris-glycine gels and transferred onto nitrocellulose. Membranes were washed with ultra-purified water and labeled with No-Stain Protein Labeling Reagent (Thermo Fisher Scientific) according to kit instructions. Next, the membranes were blocked for 45 minutes with 5% (w/v) non-fat milk in Tris-buffered saline-Tween 20 buffer (TBS-T). Primary antibodies were added to the blocking solution, and the blots were incubated overnight at 4°C with gentle agitation. Following primary incubation, blots were washed three times with 10mL of TBS-T for 10 minutes and incubated with anti-mouse and anti-rabbit fluorescent-conjugated secondary antibodies in a 1:20,000 dilution for 1 hour at room temperature. Three additional TBS-T washes with 10mL in 10 minutes were performed, and the blot signal was captured using Odyssey Fc (LI-COR).

### Antibodies

Antibodies for western blotting are used as follows: anti-*O*-GlcNAc (#ab2739-Abcam): 1:1000; Anti-actin (#A2066, Sigma Aldrich), 1:1000; Anti-Phospho-ERK1(T202/Y204)/ERK2(T185/Y187) (#AF1018, R&D systems), 1:500; Anti-ERK1/ERK2 (#AF1576,R&D systems), 1:500; Anti Phospho-p90RSK (S380) (#9341S, Cell Signaling), 1:500; Anti-Rsk1/rsk2/rsk3 (#9355T, Cell Signaling), 1:500; Anti-Phospho-Nurr77 (S351) (#5095S, Cell Signaling), 1:500; Anti-Nurr77 (#NB100-56745, Novus Biologicals), 1:500; Anti-Phospho-CREB (S133) (#9198S, Cell Signaling), 1:500; Anti-CREB (#9104S, Cell Signaling), 1:500.

### Luciferase reporter assay

AtT-20 cells were concurrently transfected using Lipofectamine 3000 (Thermofisher) with two plasmids – one containing the pRL *Renilla* luciferase gene under the control of the CMV promoter (# E2261, Promega) and the other containing the firefly luciferase gene under the control of the *Pomc* promoter (−646bp to +65bp) (#17553, Addgene). Forty-eight hours after transfection, luciferase activity was measured using the dual-luciferase assay protocol (#E1910, Promega). Transfection was performed 48h before measurement of the promoter activity.

### WGA-Pull down

*O*-GlcNAcylated Proteins were pulled down using the Triticum Vulgaris Lectin (WGA) - MagneZoom™ Kit according to the manufacturer protocol. Magnetic beads with bound proteins were washed with 200 μl of 166 mM ammonium bicarbonate. The buffer was removed using a magnetic rack, and the beads were suspended in 100 μl of 40% Invitrosol (Invitrogen), 100 mM ammonium bicarbonate. Proteins were reduced with 5 mM TCEP for 30 min at 37 °C and alkylated with 10 mM iodoacetamide for 30 minutes at 37 °C in darkness. 2 μg of trypsin/LysC mixture (Thermo Scientific Pierce) was added, and the digest was allowed to proceed overnight at 37°C. After digestion, the supernatant was removed using a magnetic rack, acidified with TFA, and desalted according to the manufacturer’s directions with the PreOmics Phoenix kit.

### Mass spectrometry

Each sample was analyzed on a Thermo Scientific Orbitrap Fusion Lumos MS via two technical replicate injections in two methods (90 or 180 min). Further details regarding these two methods can be found in supplementary methods (39).

### Pathway enrichment

Pathway enrichment was performed using the Reactome analysis tool (https://reactome.org/PathwayBrowser/#TOOL=AT) (40).

### Proliferation assay

Proliferation assays were performed using the Cell Proliferation Kit I (MTT) (#11465007001, Sigma Aldrich) according to the manufacturer’s protocol. Briefly, cells were seeded at 5×10^3^ cells well in a 96-wells plate. Each condition was performed in 6 replicates, and measurements were averaged. 10 μL of the MTT reagent was added to each well for 4 hours. 100 μL of the solubilization solution was then added overnight before measuring absorbance at 600 nm.

### Tunel assay

Apoptotic flux was measured using the EZClick™ TUNEL – in situ DNA Fragmentation/Apoptosis Assay Kit (BioVision) according to the manufacturer’s protocol. Positive control cells were treated with DNAse according to the kit’s instruction.

## RESULTS

### Clinical corticotropic tumor aggressiveness correlates with overexpression of O-GlcNAc enzymes

In all literature published on the topic, *O*-GlcNAc cycling is increased in tumors and contributes to cell-cycle regulation (26,41). Therefore, we investigated whether pituitary adenomas also overexpressed the *O*-GlcNAc cycling enzymes *OGT* and *OGA*. A pituitary gland from a recently deceased amyotrophic lateral sclerosis patient was used as a control non-tumorous pituitary tissue. For each pituitary adenoma subtypes, *OGT* and *OGA* expression levels were measured by quantitative PCR (qPCR) and normalized by ACTIN (Figure 2A/B).

**Figure 2:**
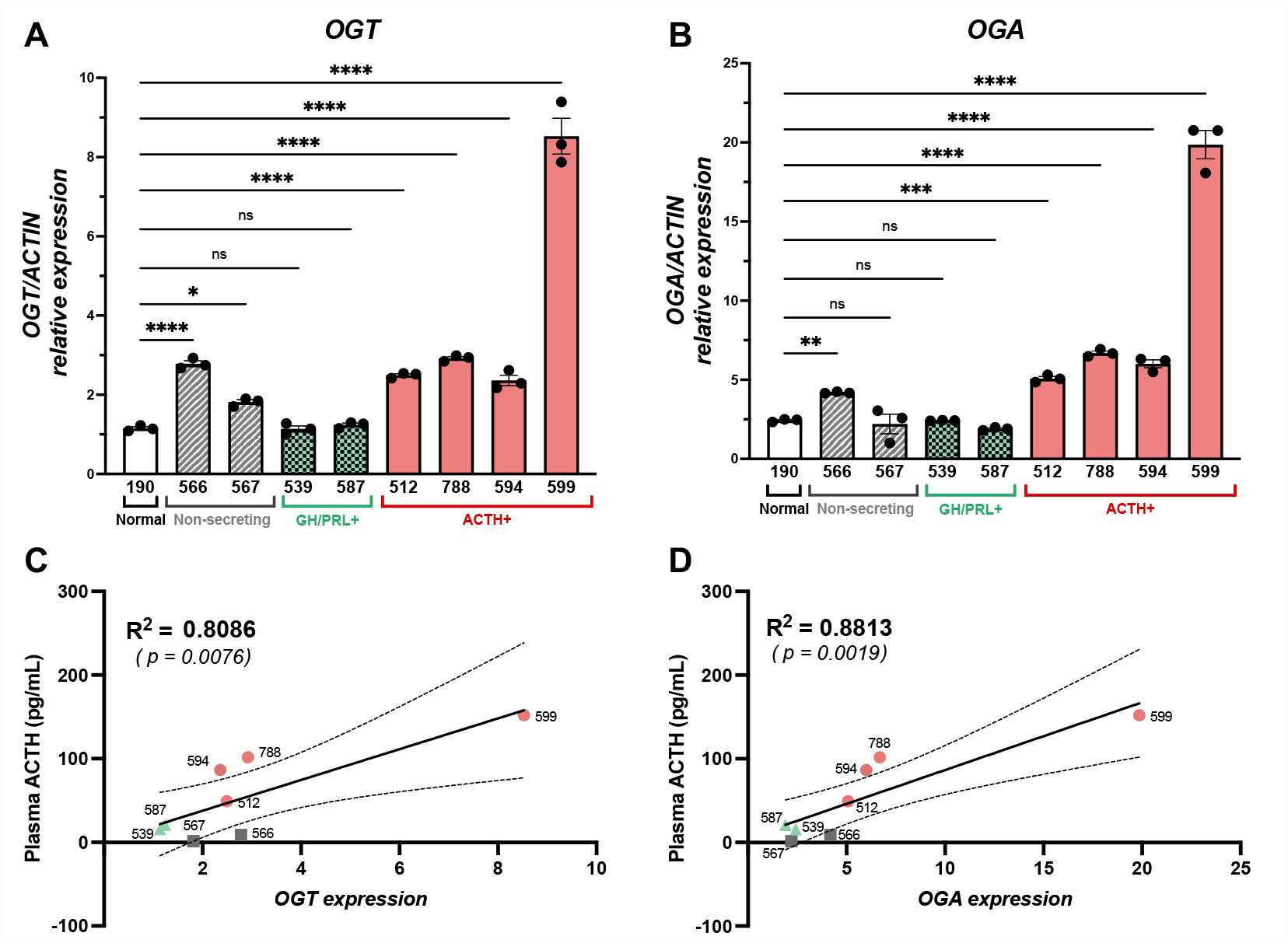
Clinical endocrine disruption in pituitary adenomas is correlated with overexpression of O-GlcNAc enzymes. **(A/B)** The expressions of *O*-GlcNAc transferase (*OGT*) (A) or *O*-GlcNAcase (*OGA*) (B) were measured by qPCR and normalized by *ACTIN*. Relative expression was calculated using the −2^ΔΔCt^ method. Significance was calculated by one-way ANOVA with Fisher’s LSD *vs*. normal pituitary tissue; *ns* ≥ *0.05*, * *p* < *0.05*, ** *p* < *0.01*, *** *p* < *0.001*, **** *p* < *0.0001*. **(C/D)** One-tailed Pearson correlations were computed between plasma ACTH and (C) *OGT* and (D) *OGA* expression (*R*^2^ = 0.81, *p* = 0.008 and *R*^2^ = 0.88, *p* = 0.002, respectively) across pituitary adenoma subtypes. Simple linear regressions with 95% confidence intervals are represented.

Non-secreting pituitary tumors showed a slight but significant upregulation of *OGT* (*n*=2), and *OGA* (*n*=1) compared to the normal pituitary gland (Figure 2A/B). On the other hand, none of the mammosomatotrophic tumors showed deregulations of *OGT* nor *OGA* (Figure 2A/B). All the corticotropic pituitary adenomas (*n*=4) exhibited a significant increase in *OGT* and *OGA* expression *vs.* control pituitary (Figure 2A/B). Interestingly, one tumor (ID #599) showed a 5-fold increase in *OGT* and *OGA* expression relative to control pituitary tissue and was clinically particularly aggressive, characterized by very high plasma ACTH and cortisol, large tumor diameter at presentation, and invasion into the clivus (Table S1) (39). Among all tumor subtypes, plasma ACTH positively correlated with OGT and OGA expression (respectively, *R^2^*=0.81, p=0.008 and *R^2^*=0.88, p=0.002, Pearson correlation), suggesting that *O*-GlcNAcylation may be involved in the hormonal function of pituitary adenomas, especially as it pertains to ACTH-secretion. In contrast, no correlation was found between *OGT* and *OGA* expression and tumor size (respectively, *R^2^*=0.38, p=0.17 and *R^2^*=0.27, p=0.26, Pearson correlation) (Figure S1) (39). Thus, we decided to focus our study on understanding the impact of *O*-GlcNAcylation on ACTH-secreting adenomas.

### O-GlcNAcylation is essential for AtT-20 cells proliferation

We first wonder if *O*-GlcNAcylation participates in the proliferation signaling in corticotropic pituitary adenomas as shown previously in other tumor environments (27,41–43). The murine AtT-20 cells were selected for this study as they possess both proliferative and endocrine abilities reminiscent of human ACTH-secreting adenomas.

Cell proliferation was measured daily by the MTT proliferation assay. Measurement of cell proliferation was measured daily by spectrophotometry in 6 replicates per condition (Figure 3A).

**Figure 3:**
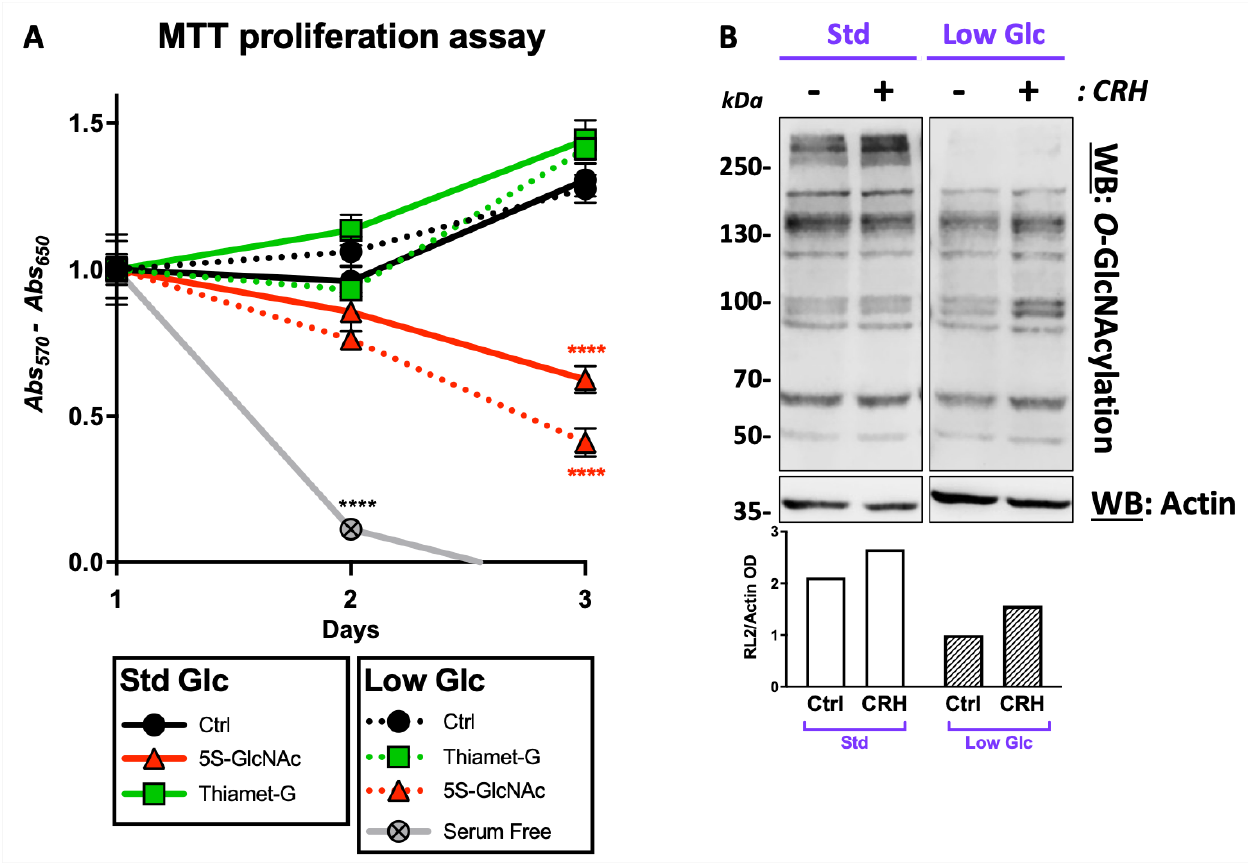
*O*-GlcNAcylation is involved in the proliferative and endocrine function of corticotropic adenomas. **(A)** Cell proliferation was measured by MTT assay in 6 replicates for each condition. Thiamet-G and 5S-GlcNAc were added daily into culture media, either standard (Std) or low glucose. Significance was analyzed by two-way ANOVA with Tukey’s post-hoc test corrected for multiple comparisons at each time point; **** *p* < *0.0001*. **(B)** AtT-20 cells were treated with CRH for four hours in standard (Std) or low glucose (Glc) media. Western blot of cell lysate with RL2 antibody probes for *O*-GlcNAcylated protein. Bottom panel represent the optical density (OD) quantification of the upper panel Western blot.

Because *O*-GlcNAcylation is a nutrient-sensing modification highly dependent on glucose level (22), AtT-20 were cultivated in both standard (Std Glc, 4.5 g/L) or low (Low Glc, 1g/L) glucose media leading to high or low basal *O*-GlcNAcylation levels (Figure 3B). As expected, switching to low glucose media lowered the overall *O*-GlcNAcylation in AtT-20 cells compared to Std media (Figure 3B). Additionally, cells were treated daily with *O*-GlcNAc transferase inhibitor 5S-GlcNAc or *O*-GlcNAcase inhibitor Thiamet-G to decrease or increase *O*-GlcNAcylation levels (Figure 3A). The efficiency of both inhibitors was confirmed by Western blot (Figure S2A/B) (39). In standard and low glucose media, 5S-GlcNAc treatment decreased cell proliferation in AtT-20 cells over the course of 3 days (Figure 3A). It was not due to increased apoptosis as measured by Tunel assay (Figure S3) (39). While the effect was moderate compared to serum-free media, cell proliferation was halved compared to the control condition (Figure 3A). Likely because their proliferative abilities are already elevated, AtT-20’s treatment with Thiamet-G did not increase proliferation and was comparable to control levels in both media conditions (Figure 3A). Overall, this emphasized the essential role of *O*-GlcNAcylation in pituitary adenomas AtT-20 cells and their potential as a druggable target to slow the expansion of ACTH-pituitary adenomas.

While extensive proliferation and expansion of pituitary adenomas is deleterious when invading neighboring structures, the first clinical signs of these tumors are often hormonal. Therefore, we next investigated the role of *O*-GlcNAcylation on the endocrine function of corticotropic pituitary adenoma cells.

### CRH stimulation increases O-GlcNAcylation levels

In AtT-20 cells – and normal corticotropes – CRH stimulation induces pro-hormone proopiomelanocortin (POMC) mRNA and protein, and finally ACTH by post-translational cleavage.

Thus, we first incubated AtT-20 cells with CRH for 4 hours to measure to levels of total *O*-GlcNAcylated protein and the expression of both *O*-GlcNAc cycling enzymes, *Ogt* and *Oga* (Figure 3B). Interestingly, CRH treatment increased overall *O*-GlcNAcylation of proteins as observed by western blot in both Std and Low glucose media (Figure 3B). Conjointly, a slight increase in *Ogt* expression was observed by qPCR in low glucose media accompanying the increase in total *O*-GlcNAcylated proteins (Figure S4A) (39). No changes in *Oga* were noted (Figure S4B) (39).

This suggested that CRH promoted a quick rise in *O*-GlcNAcylated proteins, likely on signaling molecules, but did not dramatically alter *O*-GlcNAc enzymes’ expression. Thus, we next examined the impact of *O*-GlcNAcylation on CRH-downstream signaling.

### O-GlcNAcylation stimulates CRH downstream signaling

CRH stimulates downstream signaling in corticotropes, including the MAP kinase signaling cascade, ultimately resulting in *Pomc* transcription (Figure 4A).

**Figure 4:**
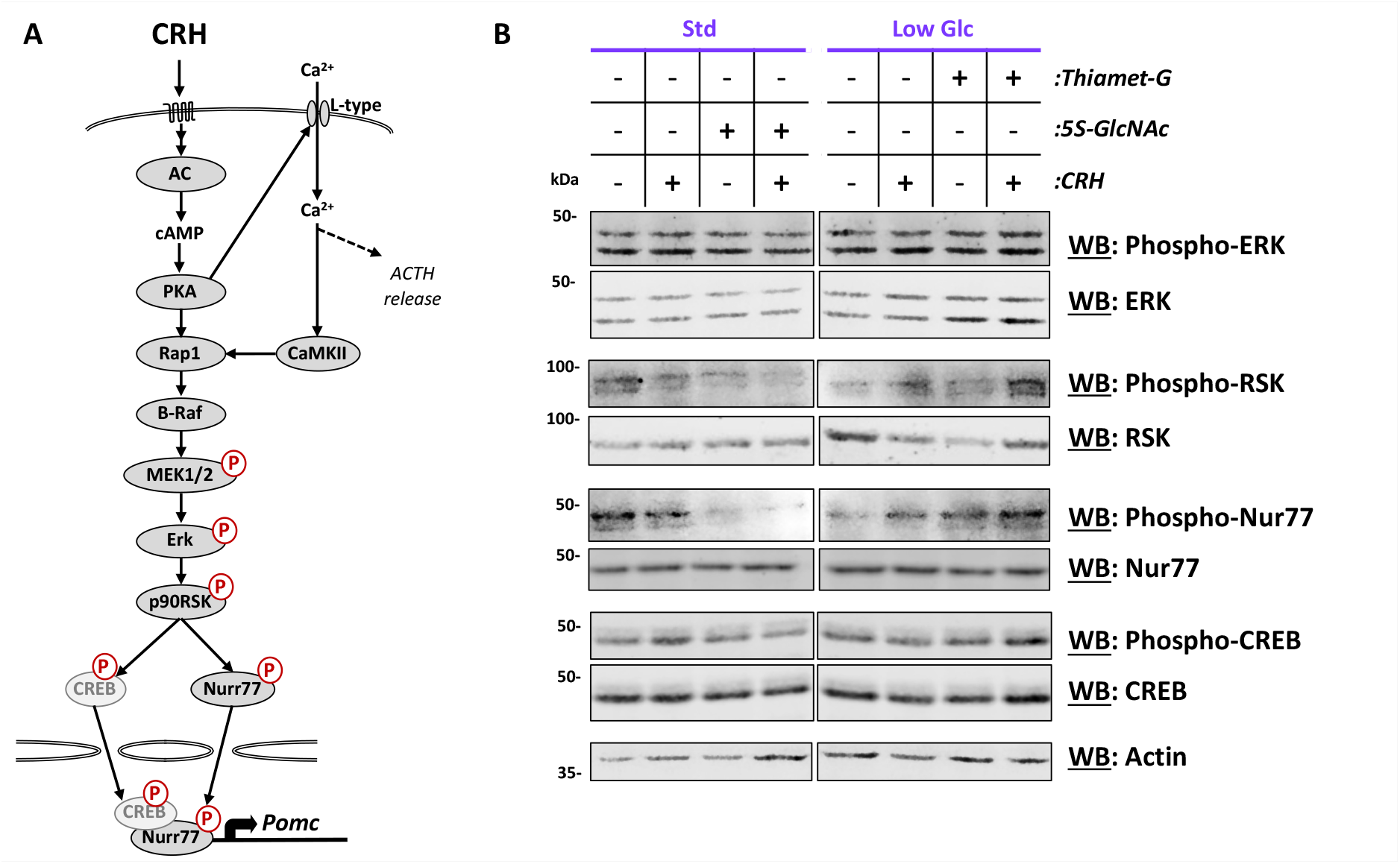
*O*-GlcNAcylation is necessary for downstream MAPK signaling pathway phosphorylation. **(A)** Schematic of downstream targets of CRH-trigger signaling pathway in corticotropes. **(B)** AtT-20 cells were pretreated overnight with 5S-GlcNAc or Thiamet-G. Levels of phosphorylated and total protein for several downstream MAPK targets were measured by western blot after CRH treatment (4h).

In both standard and low glucose media, AtT-20 cells were treated with CRH for 4 hours. Analysis of the MAPK effectors and their activated/phosphorylated forms was performed by Western blot. In addition, AtT-20 cells in standard glucose media were also pre-treated with 5S-GlcNAc to prevent *O*-GlcNAcylation (Figure S2A) (39). In low glucose/*O*-GlcNAcylation condition, AtT-20 were also pre-treated with Thiamet-G to increase total *O*-GlcNAcylation, independently of glucose availability (Figure S2B) (39). In Std glucose media, activation of the MAPK signaling pathway following CRH treatment has been shown to happen within 30 min (44,45) (Figure S5) (39). Therefore, MAPK effector phosphorylation was minimal at the 4h time point in CRH-stimulated *vs*. Ctrl cells (Figure 4B). However, CRH-stimulation of the MAPK signaling pathway was still visible in the low glucose condition at 4h post-CRH, including increased phospho-RSK and phospho-NURR77, representing slower signaling (Figure 4B). However, in both unstimulated and CRH-treated AtT-20 cells, phospho-RSK and phosphor-NURR77 were dramatically reduced when pre-treated with the *O*-GlcNAc inhibitor 5S-GlcNAc (Figure 4B). It emphasized a role for *O*-GlcNAcylation in regulating basal and stimulated MAPK signaling in corticotropic pituitary adenomas.

Finally, increasing *O*-GlcNAcylation levels with Thiamet-G in low glucose media increased basal and CRH-stimulated phospho-RSK and phospho-NURR77(Figure 4B), confirming the critical role of *O*-GlcNAc in MAPK basal activation.

We next interrogated whether alterations in MAPK signaling dependent on *O*-GlcNAcylation levels also impact *Pomc* mRNA, the ultimate product of the CRH signaling pathway.

### Total Pomc mRNA availability is dependent on O-GlcNAcylation level

At 4h post-CRH stimulation, *Pomc* expression was measured by RT-qPCR and normalized by *Actin* (Figure 5A/B). In Std media, CRH induced *Pomc* expression (Figure 5A). This was prevented by 5S-GlcNAc-mediated inhibition of *O*-GlcNAcylation, as well as by another OGT inhibitor, OSMI4 (Figure 5A, S2A/C) (39). Furthermore, low glucose/*O*-GlcNAc conditions suppressed the ability of CRH to induce *Pomc* expression (Figure 5B). However, this aptitude was recovered by supplementing media with Thiamet-G or by transfection of siRNA against *Oga* and boosting proteins’ *O*-GlcNAcylation (Figure 5B, S2B/D, S6) (39).

**Figure 5:**
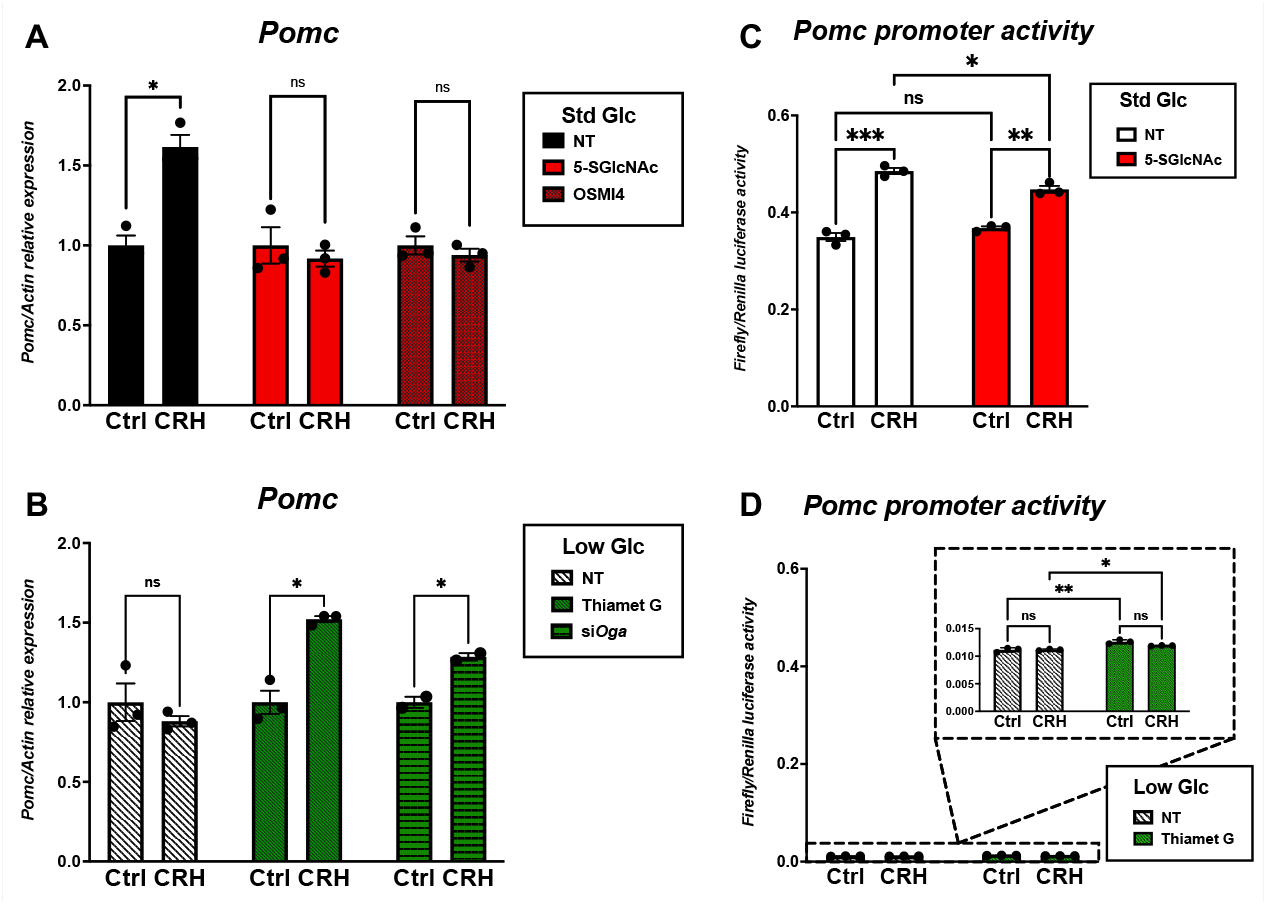
O-GlcNAcylation impacts total *Pomc* mRNA level but does not influence *Pomc* promoter activity (A/B). Proopiomelanocortin (*Pomc*) expression was measured after CRH treatment (4h) in either standard (Std) (A) or low glucose conditions (B). Cells were pre-treated overnight with 5SGlcNAc, OSMI4, Thiamet-G or transfected with siRNA against Oga. Significance was analyzed by two-way ANOVA with Fisher’s LSD; *ns* ≥ *0.05*, * *p* < *0.05*, ** p < *0.01*. **(C/D)** Pomc promoter activity was measured by firefly luciferase and reported on Renilla luciferase after CRH treatment (4h) in **(C)** standard (Std) or **(D)** low glucose conditions. Cells were pretreated overnight with 5S-GlcNAc or Thiamet-G. Significance was calculated by two-way ANOVA with Fisher’s LSD; *ns* ≥ *0.05*, * p < *0.05*, ** p < *0.01*, *** p < *0.001*.

This experiment demonstrated that sufficient glucose level, and subsequent *O*-GlcNAcylation, are critical to promoting *Pomc* expression upon CRH stimulation, which we presumed was due to enhanced *Pomc* Promoter activity.

### O-GlcNAcylation has a limited effect on Pomc promoter activity

A *Pomc* luciferase reporter assay was used to measure the *Pomc* promoter’s activation Cells were co-transfected with two constructs: *PGL3-Pomc-Luc,* expressing the firefly luciferase under the control of the *Pomc* promoter and *pRL*, expressing the Renilla luciferase under a general mammalian promoter. Std and low glucose conditions and *O*-GlcNAc inhibitors were used as previously described, in addition to CRH stimulation. Firefly luciferase activity was measured by luminescence and reported on Renilla used as the internal control for transfection efficiency (Figure S7).

Agreeing with total *Pomc* mRNA levels, CRH induced *Pomc* promoter activity in standard but not low glucose condition, demonstrated respectively by an increase or no increase in firefly luciferase activity (Figure 5C/D). However, 5S-GlcNAc only mildly suppressed this stimulation (Figure 5C). Simultaneously, Thiamet-G only slightly increased the basal promoter activation but did not rescue low glucose levels (Figure 5D). This is in divergence with the effect of Thiamet-G on total *Pomc* mRNA levels. However, *O*-GlcNAcylation seemed to slightly increase the unstimulated basal level of *Pomc* promoter activation (Figure 5D).

Overall, this suggested that glucose but not *O*-GlcNAcylation was critical for CRH-dependent *Pomc* promoter stimulation. However, because this experiment did not recapitulate the finding that *O*-GlcNAcylation is essential to increase *Pomc* expression upon CRH stimulation, we next searched for alternative pathways stimulated by CRH that would impact *Pomc* mRNA in an *O*-GlcNAc-dependent fashion.

### O-GlcNAc-dependent mRNA processing pathways are impacted by CRH treatment

*O*-GlcNAcylated proteins from AtT-20 cells treated or untreated with CRH for 1h were pulled down using the Wheat Germ Agglutinin (WGA) lectin. WGA-bound proteins were then analyzed by EThcD-MS to detect *O*-GlcNAcylated residues and quantify the relative changes in *O*-GlcNAcylated proteins upon CRH treatment (Figure 6A/B, Table S2/3) (39). A total of 1051 and 986 proteins were identified and mapped to the mouse proteome in control and CRH-treated AtT-20 cells, respectively.

**Figure 6:**
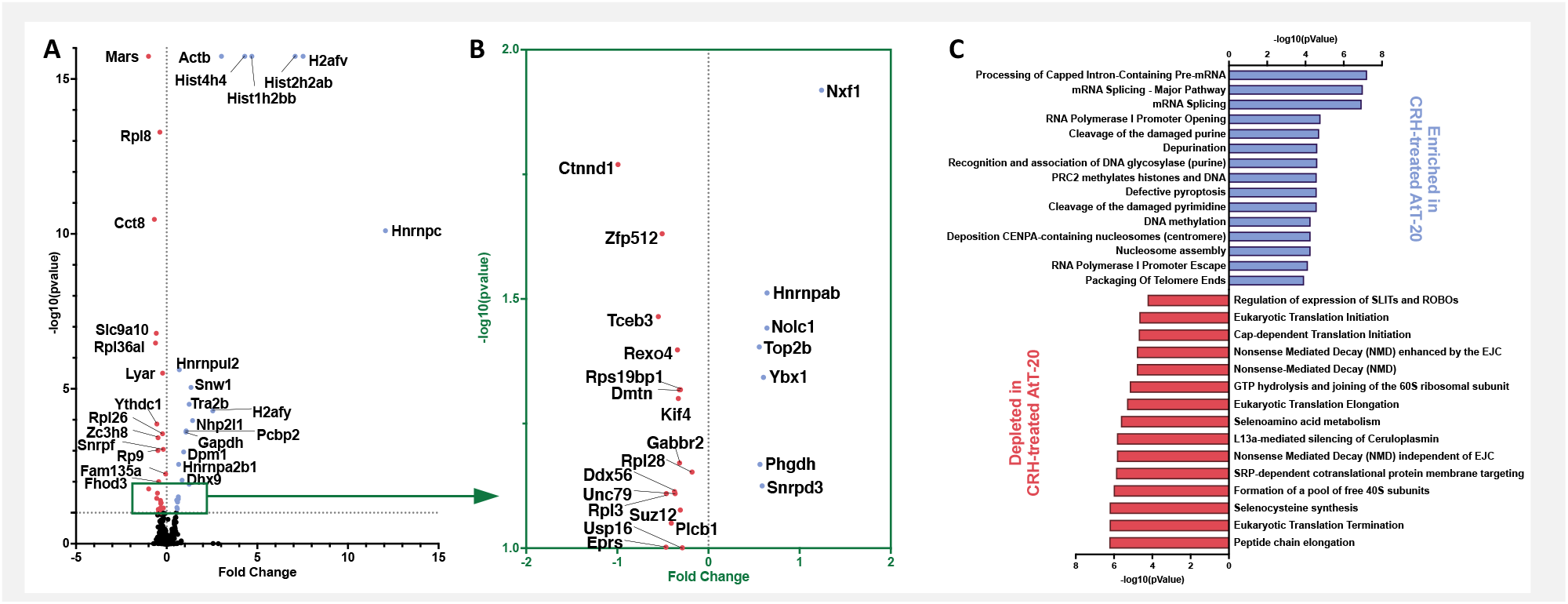
CRH induces O-GlcNAc-dependent changes in mRNA processing. **(A/B)** Fold change of *O*-GlcNAcylated protein pulled down with Wheat Germ Agglutinin (WGA) following CRH stimulation (1h) in AtT-20 cells. **(B)** Magnified view of (A) between - log_10_(pvalue) of 1 to 2. **(C)** Pathway enrichment analysis for *O*-GlcNAcylated proteins enriched or depleted in CRH-treated AtT-20 cells.

Amongst these, 399 *O*-GlcNAc sites were identified on 321 proteins (Table S2/3) (39). More than half of the proteins identified have already been identified in previous *O*-GlcNAcomics experiments (46), confirming the validity of our experiments (Figure S8, Table S4) (39). Measurement of peak area for relative quantification highlighted 52 proteins differentially pulled down by WGA in control and CRH-treated AtT-20 cells (Figure 6A/B, Table S5) (39). Significantly deregulated proteins were inputted into the analysis tool of the Reactome platform (40) for enrichment analysis (Figure 6C, Table S6/7) (39). Enrichment of “mRNA processing” pathways, including “processing of intron-containing pre-mRNA” and “mRNA splicing,” “transcription,” and “DNA repair,” was noted in CRH-treated cells (Figure 6C, Table S6) (39). Depleted in CRH-treated cells (versus control) were protein processing pathways like “translation,n” and “non-sense mediated decay (NMD),” a central pathway for mRNA degradation (Figure 6C, Table S7) (39).

This analysis demonstrated that *O*-GlcNAcylations of pre-mRNA splicing and mRNA stability factors are modulated by CRH, identifying these pathways as potential mechanisms for the regulation of *Pomc* expression.

### Pomc mRNA stability is regulated by O-GlcNAcylation

To investigate whether *O*-GlcNAcylation participates in *Pomc* mRNA stabilization, Pomc expression was measured in AtT-20 cells treated with the mRNA synthesis DRB. An 8h time-course measuring *Pomc* expression normalized by *Actin* by qPCR confirmed the rapid decay of *Pomc* mRNA in less than 2h. (Figure S9A) (39). While the basal level of *Pomc* expression was reduced by two-fold in low glucose levels, DRB was still effective in preventing the neo-synthesis of *Pomc* mRNA (Figure S9B) (39), confirming the compatibility with our previous experimental design. Concurrent treatment of DRB with CRH prevented *Pomc* mRNA decay in standard and low glucose levels showing, for the first time, that CRH rapidly stabilizes *Pomc* mRNA (Figure 7A). However, The effect was more permanent in Std glucose condition and persisted two hours after DRB treatment (Figure 7A). Thus, per the *O*-GlcNAcomic analysis, we demonstrated that, while *Pomc* mRNA is subject to rapid decay, CRH stimulation prevented *Pomc* mRNA degradation. We next investigated whether *O*-GlcNAcylation affects this process by concurrently treating AtT-20 cells with OSMI4 or Thiamet-G to decrease and increase *O*-GlcNAcylation levels (Figure S2B/C/D/E) (39). Following overnight OSMI4 treatment, *Pomc* mRNA decay was increased regardless of CRH treatment (Figure S2B/C) (39). A similar response was observed following 5S-GlcNAc treatment, another OGT inhibitor (Figure S9C) (39). The fact that OSMI4 also affected *Pomc* mRNA decay in Ctrl condition suggested that overnight treatment with *O*SMI4 or 5S-GlcNAC likely downregulated the expression of critical mRNA regulatory protein as previously published (47–50). Therefore, we repeated the experiment by adding OSMI4 only 1h before DRB treatment, thus only affecting the ability to cycle *O*-GlcNAcylation onto existing protein (Figure S9D/E) (39). While OSMI4 did not impact *Pomc* decay in Ctrl condition, OSMI4 prevented the stabilization of *Pomc* mRNA in CRH-treated cells (Figure S9E) (39), suggesting that rapid *O*-GlcNAc cycling is regulating *Pomc* mRNA turnover upon CRH stimulation. Inversely, in low glucose, co-treating cells with Thiamet-G and CRH stabilized *Pomc* mRNA compared to CRH alone (Figure 7D/E), suggesting that *O*-GlcNAcylation is required for CRH-induced *Pomc* mRNA stabilization.

**Figure 7:**
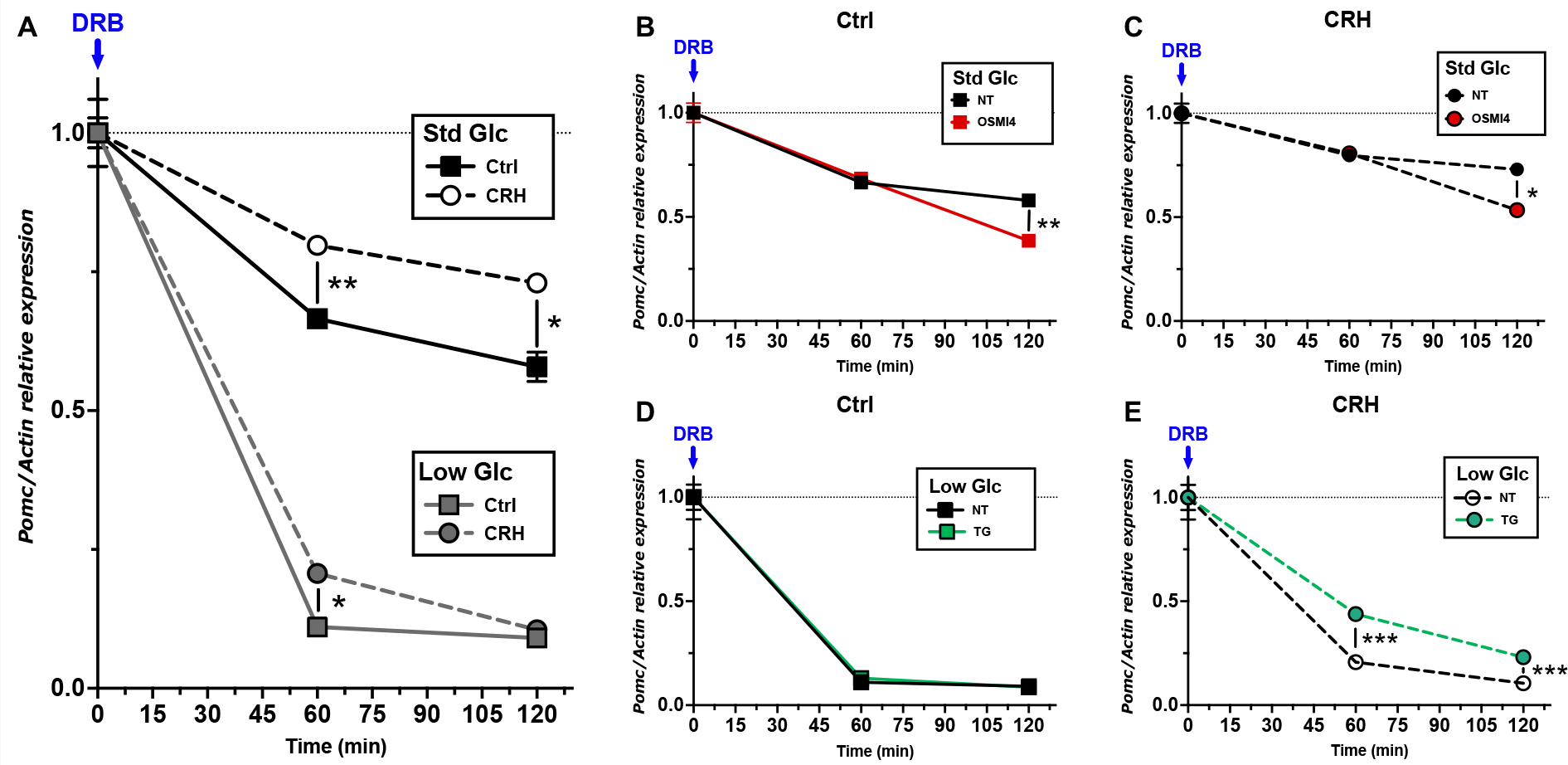
CRH stabilizes Pomc mRNA in an O-GlcNAc-dependent manner. **(A)** Proopiomelanocortin (*Pomc*) mRNA level was measured by qPCR after cotreatment with CRH and the transcription inhibitor 5,6-dichloro-1-beta-D-ribofuranosysl-benzimidazole (DRB) for 2h in standard (Std) or low glucose conditions. **(B/C/D/E)** Cells were pre-treated overnight with OSMI4 **(B/C)** or Thiamet-G **(D/E)**. Significance was measured by two-way ANOVA with Fisher’s LSD; *ns* ≥ *0.05*, ** *p* < *0.01*, *** *p* < *0.001*, **** *p* < 0.0001

Overall, this experiment stressed the essential contribution of *O*-GlcNAcylation in CRH-mediated *Pomc* expression, specifically by promoting the stabilization of *Pomc* mRNA. Interestingly, improper splicing of mRNA is one of the mechanisms leading to mRNA decay and thus impacting mRNA’s half-life. Interestingly, mRNA splicing was also highlighted as enriched in CRH-stimulated cells as presented above (Figure 6C, Table S6) (39). Thus, we next investigated the possibility that *O*-GlcNAcylation and CRH stabilize *Pomc* mRNA by promoting proper splicing.

### Intron-containing mRNA levels are dependent on O-GlcNAcylation

Since *O*-GlcNAcomic analysis showed that “processing of intron-containing pre-mRNA” was the most enriched *O*-GlcNAc-dependent pathway upon CRH stimulation (Figure 6C, Table S6) (39), we investigated the impact of *O*-GlcNAcylation on intron-containing *Pomc* RNA. A specific set of primers spanning the last intron into the last exon was used to measure the level of intron-containing *Pomc mRNA* by PCR (Figure 8A). Inhibition of OGT by OSMI4 led to increased intron-containing *Pomc* mRNA in both control and CRH-treated cells. OSMI4 also lead to the appearance of shorter PCR products, likely corresponding to *Pomc* mRNA species with partially retained introns (Figure 8B). In low glucose conditions, increasing quantities of partially retained intron species of *Pomc* mRNA were noted compared to Std glucose condition. This correlated with the limited ability of CRH to induce *Pomc* mRNA stabilization in low glucose levels (Figure 8B). On the other hand, when treated with Thiamet-G, intron-containing *Pomc* mRNA species were dramatically reduced, potentially due to enhanced proficiency in splicing (Figure 8B).

**Figure 8:**
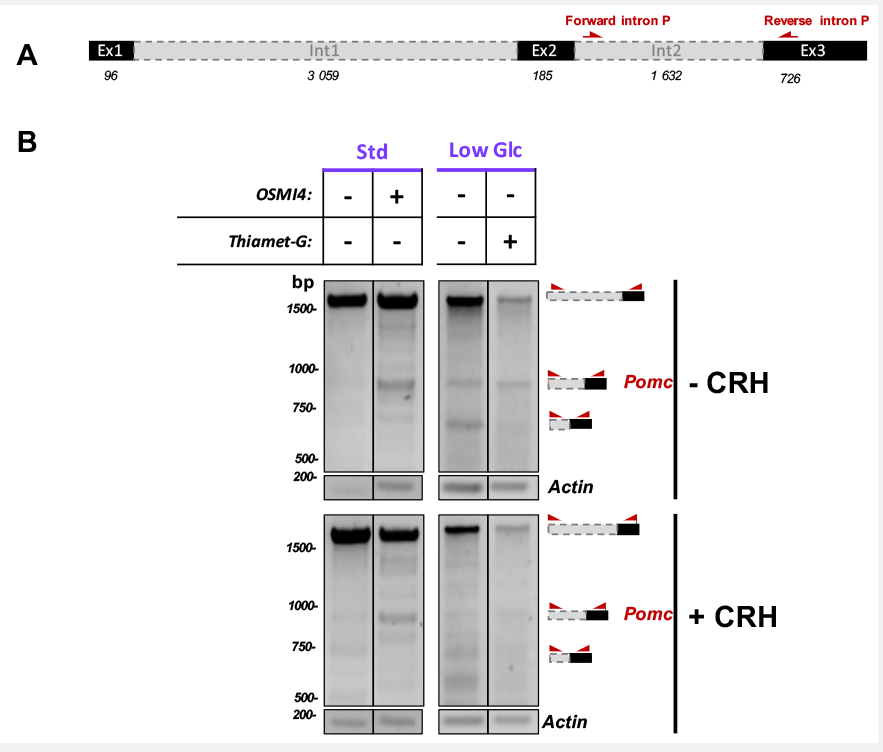
*O*-GlcNAcylation reduces the levels of introncontaining Pomc mRNA. **(A)** Primers used for measure intron containing Pomc mRNA. **(B)** RT-PCR was performed on CRH-treated (4h) or Ctrl AtT-20 cells total mRNA pre-treated with OSMI4 or Thiamet-G overnight in standard (Std) or low glucose conditions.

Taken together, we showed that *O*-GlcNAcylation regulates *Pomc* mRNA intron splicing, hypothetically by facilitating proper mRNA splicing, contributed to *Pomc* stabilization by CRH.

### Bioinformatic analysis reveals *O*-GlcNAcylated targets involved in regulation of mRNA processing

To identify *O*-GlcNAcylated proteins responsive to CRH and potentially responsible for regulating Pomc mRNA, we explored the *O*-GlcNAcylated proteins identified previously by WGA pull-down and mass spectrometry (Figure 6A/B, Table S2/3) (39). In AtT-20 cells, 321 *O*-GlcNAcylated proteins and 399 *O*-GlcNAcylation sites were identified (Table S4) (39). For those, we extracted information on functional domains and functions from UniprotKB and linked databases. Thus, 24 *O*-GlcNAcylated proteins were determined to be involved in mRNA binding and/or processing and are excellent targets for CRH-mediated regulation of *Pomc* mRNA. Additionally, 180 and 81 proteins carried phosphoserine and phosphothreonine sites, respectively (Figure 9A, Table S8) (39), and might be the subject of *O*-GlcNAcylation/ phosphorylation interplay. Therefore, we provide a condensed list of 24 *O*-GlcNAcylated proteins in pituitary adenomas cells that are likely involved in *Pomc* mRNA regulation upon CRH stimulation (Figure 9A).

**Figure 9:**
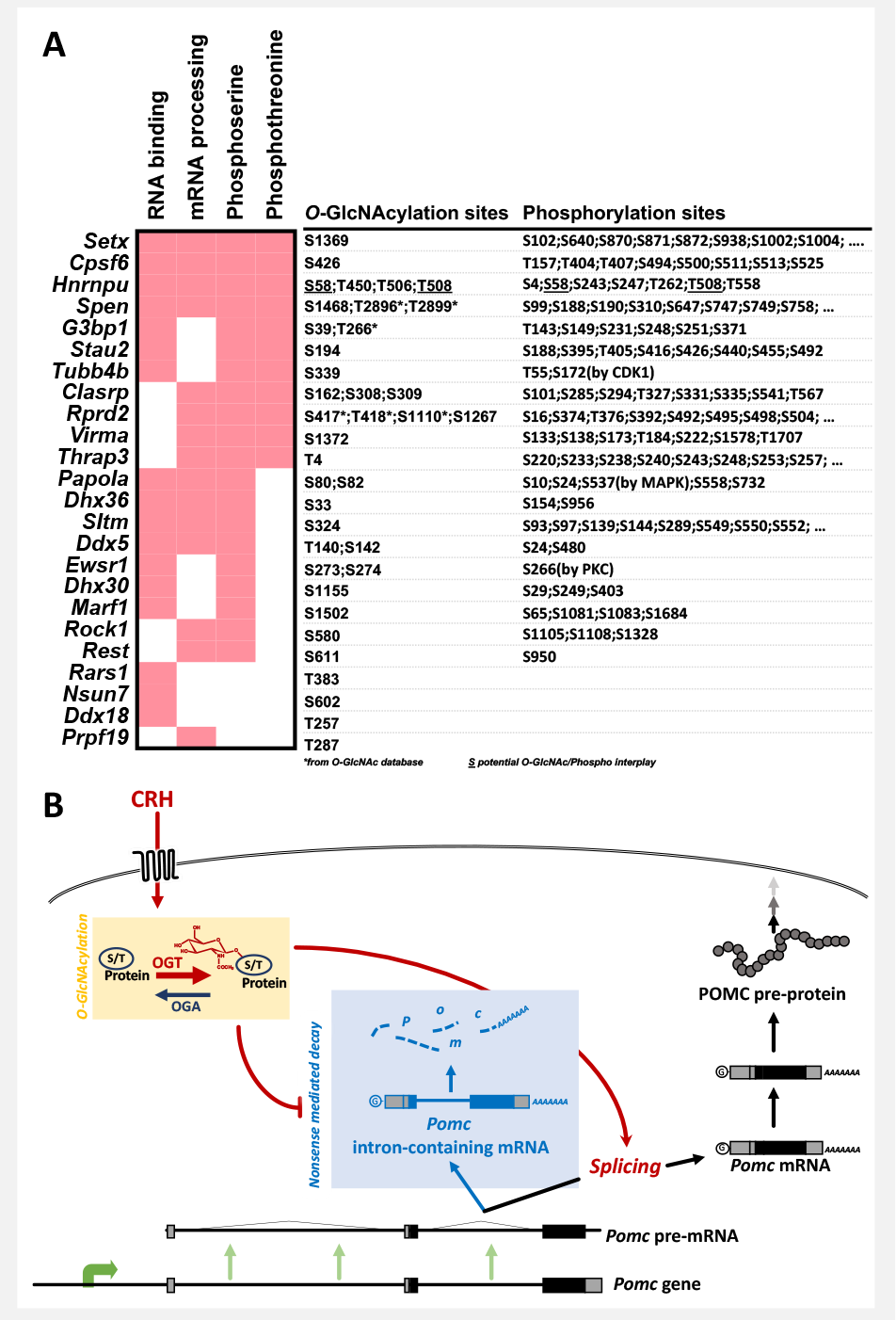
Bioinformatic analysis reveals potential targets for regulation of *Pomc* mRNA stability. **(A)** List of *O*-GlcNAcylated targets identified as potential regulators of *Pomc* mRNA stabilization. Phosphorylation status and mRNA binding ability as well as their involvement in mRNA processing were extracted from UniprotKB (https://www.uniprot.org). Additional O-GlcNAcylated sites (*) were extracted from the O-GlcNAc database (https://www.oglcnac.mcw.edu). **(B)** AtT-20 cells transcribe and degrade *Pomc* mRNA in a steady-state balance in the absence of CRH. When stimulated by CRH, existing *Pomc* mRNA is stabilized, allowing for rapid expression of *Pomc*. O-GlcNAcylation promotes proper intron-splicing to stabilize *Pomc* mRNA upon CRH stimulation.

## DISCUSSION

Effective medical treatment of Cushing’s disease, defined as a hypercortisolism due to a corticotropic pituitary adenoma, has so far been limited to systemic endocrine modulation such as glucocorticoid receptor (GR) blockade and dexamethasone suppression. Meanwhile, the treatment of prolactinomas was revolutionized by the revelation that dopamine agonists can counter the hypersecretion of prolactin (PRL) and cause tumor regression in many cases (51). Likely because of this success, dopamine agonists have been piloted to treat other subtypes of pituitary adenoma. Clinical studies of these drugs have not demonstrated satisfactory efficacy in treating non-PRL pituitary adenomas (52). Therefore, it is imperative to better understand the unique cell biology of corticotropic pituitary adenomas and further our understanding of these tumors to guide drug discovery.

Pituitary adenomas have been studied through many different lenses, including such broad studies as epigenomics, mitochondrial genomics, and gene expression microarrays. These screening studies have highlighted many dysregulated pathways in various pituitary adenoma subtypes, including those involved in TGF-beta signaling, colorectal cancer, amino and nucleotide sugar metabolism, and *O*-glycan biosynthesis (53). Aberrations in the nutrient-sensitive *O*-GlcNAc posttranslational protein modification have been implicated in a vast array of tumor types (27). While this posttranslational protein modification competes for phosphorylation sites, *O*-GlcNAcylation is regulated by only two enzymes rather than various kinases and phosphatases. OGT and OGA are responsible for cycling *O*-GlcNAc moieties at serine/threonine residues, thereby modulating countless biochemical processes from transcription to RNA processing to cell cycle regulation (21). The extensive involvement of *O*-GlcNAc cycling in tumor biochemistry and the existence of numerous enzyme inhibitors targeting *O*-GlcNAc cycling make it a compelling target for therapeutics. In pituitary corticotropes, OGT has been shown to be involved in GR-mediated transcriptional repression (54). Further, glucose uptake appears to be upregulated at the cell membrane in response to CRH, which would provide an additional precursor for *O*-GlcNAc (30). It is, therefore, essential to understand the regulation of the *O*-GlcNAc cycling enzymes OGT and OGA in the context of pituitary adenomas.

Our study demonstrates that *OGT* and *OGA* are preferentially upregulated in corticotropic pituitary adenomas (and, to a lesser extent, non-secreting pituitary adenomas) (Figure 2). Furthermore, we demonstrate that the degree of upregulation of *OGT* and *OGA* correlates well with the degree of hypersecretion of ACTH observed clinically. This indicates a potential role for *O*-GlcNAc cycling in the aberrant regulation of ACTH secretion in pituitary adenomas. Furthermore, we show a specific case of a particularly young patient with an aggressive invasive corticotrope pituitary adenoma and marked associated hypercortisolism. We, therefore, speculate that *O*-GlcNAc enzymes may be a useful pathological biomarker for identifying invasive potential in pituitary adenomas and certainly seems to be a reliable marker of profound endocrine disruption.

### Corticotropic pituitary adenoma proliferation is O-GlcNAcylation-dependent

In numerous cancers, *O*-GlcNAcylation and/or *O-*GlcNAc enzymes are dysregulated (26,41). Unsurprisingly then, many oncogenes and tumor suppressors are *O*-GlcNAcylated, including Ki-67 (55), β-catenin (56), E-cadherin (57), galectin-3 (58), and p53 (59), all of which are markers of pituitary adenoma invasiveness (60). For some of these proteins, *O*-GlcNAcylation’s role has been well defined. For example, the *O*-GlcNAcylation of β-catenin stabilizes the protein and drives its transcriptional activity, resulting in increased proliferation (56,61). Similarly, p53 *O-*GlcNAcylation affects its degradation in cancer cells (59,62). Therefore, it is not surprising that preventing *O-*GlcNAcylation would affect a network of *O-*GlcNAcylated oncogenes, ultimately resulting in suppressed pituitary adenoma proliferation *in vitro* (Figure 3A). Consequently, we are suggesting that *O*-GlcNAcylation might be both a diagnostic tool and a druggable to reduce pituitary adenoma progression (63). After establishing the importance of *O*-GlcNAcylation in regulating corticotropic adenoma proliferation, we addressed the similarly important issue of hormone expression regulation in these cells.

### O-GlcNAc cycling participates in CRH-dependent signaling pathway

In stressful situations, circadian stimulation, and other factors, the hypothalamus releases CRH into the hypophyseal portal circulation, which carries it to the target pituitary corticotropes, inducing them to synthesize, and secrete ACTH. To do so, the binding of CRH to its receptor triggers a signaling cascade, including both protein kinase A (PKA) and Ca^2+^/Calmodulin-dependent protein kinase II (CAMKII)-dependent phosphorylation cascade leading to neo-synthesis of *Pomc* mRNA and protein (64,65) (Figure 4A). Interestingly, stimulating pituitary adenoma cells with CRH leads to an overall increase in *O*-GlcNAcylated protein, demonstrating that *O*-GlcNAcylation is responsive to CRH stimulation (Figure 3B).

A relationship between *O*-GlcNAcylation and intracellular signaling was described shortly after its discovery. Due to its interplay with serine and threonine phosphorylation, *O*-GlcNAc often counteracts phosphorylation-driven signaling, including interactions with insulin and various MAP kinase signaling pathways (66–70). For example, OGT attenuates insulin signaling by *O*-GlcNAcylation of proteins involved in numerous steps in the PI3 kinase signaling pathway (71). In addition, *O*-GlcNAcylation also impacts p38 MAPK pathways by modifying JNK (c-Jun N-terminal kinase) and ASK1 (apoptosis signal-regulating kinase 1) (67). Finally, in gastric cancer, elevated *O*-GlcNAcylation promotes ERK 1/2 signaling and cell proliferation (68), linking *O*-GlcNAcylation to extracellular signal-dependent MAPK/ERK signaling, which is central to the CRH-responsiveness of corticotropes.

Of the proteins involved in CRH signal transduction, many are *O*-GlcNAcylated (21), including PKA (72), CAMKII (73), B-RAF (74), MEK1 (74), extracellular signal-regulated kinase 1/2 (ERK1/2) (75), RSK (76,77), CREB (78) and NURR77 (79) (Figure 4A). Therefore, we anticipated that *O*-GlcNAc modulation would affect the ability of CRH to induce downstream signaling. Not surprisingly, preventing *O*-GlcNAcylation reduced the basal activation of several MAPK effector proteins in AtT-20 cells (Figure 4B). While little *O*-GlcNAcylation suppressed basal MAPK activation, we found no evidence of an interaction with CRH. Therefore, we concluded that *O*-GlcNAcylation increases the basal activation of the MAPK pathway in corticotropic cells, independently of CRH stimulation. In pituitary tumors, this is particularly relevant as CRH stimulus is usually not required for increased ACTH production (80). It also agrees with our observation that human ACTH+ pituitary adenomas demonstrate increased expression of *O*-GlcNAc cycling enzymes, and these increases correlate well with plasma cortisol levels (Figure 2).

While these observations indicate an intricate relationship between *O*-GlcNAcylation and corticotrope function, the lack of interaction between CRH and *O*-GlcNAcylation in MAPK activation suggests *O*-GlcNAcylation impacts signaling downstream of the MAPK cascade.

### O-GlcNAcylation increases available Pomc mRNA level by stabilizing an existing pool of mRNA

One of the downstream products CRH/MAPK pathway is the neo-synthesis of *Pomc* mRNA and, ultimately, secretion of ACTH. We first demonstrated that *O*-GlcNAcylation was essential for that outcome. Indeed, preventing *O*-GlcNAcylation precluded CRH-induction of *Pomc* expression (Figure 5A). Since *O*-GlcNAcylation is highly glucose-dependent, lowering the glucose level in AtT-20 cells prevented CRH-induced *Pomc* expression, but we could bypass this inhibition by artificially increasing *O*-GlcNAcylation (Figure 5B). These data indicate that *O*-GlcNAcylation is necessary for CRH induction of *Pomc* expression, so we were surprised to find that *O*-GlcNAcylation only mildly affected *Pomc* promoter activation (Figure 5C/D). We, therefore, concluded that the critical regulatory step whereby CRH and *O*-GlcNAcylation interact must also be post-transcriptional.

Using an *O*-GlcNAcomic approach and pathway enrichment analysis, we discovered that stimulation by CRH induces changes in *O*-GlcNAcylated proteins participating in mRNA processing. Specifically, we observed increased levels of *O*-GlcNAcylated splicing regulators and decreased levels of *O*-GlcNAcylated mRNA decay proteins (Figure 6). Over recent years, a relationship between *O*-GlcNAc and mRNA metabolism has been demonstrated. In a meta-analysis of the *O-* GlcNAcome, we previously highlighted that the most enriched ontology of the human *O-* GlcNAcome was RNA processing, including various mRNA decay mechanisms and binding proteins (21). Many mRNA surveillance proteins are *O*-GlcNAcylated, including AUF1 (55,81), ZFP36-like-1 (82), KSRP (55,83), RHAU (81), CUG-BP1 (55,84), and CUG-BP2 (84). Nevertheless, the function of their *O*-GlcNAcylation remains to be characterized. Based on the existing literature and the present *O*-GlcNAcomic analysis, we imagine that CRH, through downstream signaling, might affect the *O*-GlcNAcylation of mRNA processing elements to regulate *Pomc* mRNA levels.

In *Xenopus* melanotropes, it was previously suggested that *Pomc* mRNA is post-transcriptionally regulated via an mRNA-degradation system (37). While in the context of light adaptation, this study concluded that *Pomc* mRNA induction did not require neo-synthesis and led us to look deeper into *Pomc* mRNA decay. To this end, we demonstrate that an existing pool of *Pomc* mRNA is subjected to a rapid decay in unstimulated cells. Furthermore, we demonstrate, for the first time, that CRH induction can mitigate this decay, quickly resulting in a stabilization of *Pomc* mRNA in an *O*-GlcNAc-dependent manner (Figure 7).

While mRNA stabilization mechanisms have been studied for other pituitary hormones, the regulatory proteins involved in *Pomc* mRNA degradation remain unknown (31–33,35,36,85–87). In light of our *O*-GlcNAcomic analysis, intron retention and associated degradation by nonsense-mediated decay appeared to be the most plausible mechanism for this rapid change in *Pomc* mRNA stability. Indeed, we demonstrated intron retention in *Pomc* mRNA, which is modulated by changes in *O*-GlcNAcylation (Figure 8/9B). Not surprisingly, intron retention has been previously demonstrated to be regulated by O-GlcNAcylation (50).

The specific factors that must be *O*-GlcNAcylated to enhance splicing and thus stabilize the transcribed pool of *Pomc* mRNA are likely numerous given the complexity of splicing and the relative lack of specificity of the *O*-GlcNAc cycling enzymes. By integrating our *O*-GlcNAcomic analysis with data obtained from publicly available protein databases, we highlighted more than 20 *O*-GlcNAcylated proteins that are excellent candidates to regulated *Pomc* mRNA in an *O*-GlcNAc- and CRH-dependent manner. First, hnRNPU, a critical splicing factor, contains two known sites, both *O*-GlcNAcylated and phosphorylated, indicating significant regulatory potential that might be involved in direct competition between those two modifications in response to CRH (Figure 9A). In a previous study, hnRNPU was shown to be one of the myriad proteins phosphorylated by PKA, central to intracellular signal transduction in response to CRH stimulation (88). Another protein identified in our analysis, *Papola*, is phosphorylated and regulated by Erk and was found to be *O*-GlcNAcylated in our experiments, indicating another protein with significant potential for crosstalk between diverse signaling pathways (89) (Figure 9A). A deeper investigation of the direct competition between phosphorylation and *O*-GlcNAcylation on these targets, as well as their binding to mRNA, will be necessary to define the mechanism by which CRH stabilizes *Pomc* mRNA.

In conclusion, this study uncovered the importance of *O*-GlcNAcylation in regulating proliferation and endocrine function in corticotropic adenomas. While this study emphasized the role of *O-*GlcNAcylation in regulating *Pomc* mRNA turnover and the responsiveness to CRH stimulation, future work will be needed to demonstrate the clinical implications of this novel regulatory mechanism and identify potential sites for pharmacologic manipulation of this pathway.

## ACKNOWLEDGMENTS

We first thank Dr. Peter LaViolette, Allison Lowman, and Dr. Mona Al-Gizawiy (MCW) for supporting pituitary tissue collection and banking. Furthermore, we thank Dr. Hershel Raff (MCW) for his guidance with this project. We also thank Drs. Anat Ben-Shlomo and Melmed Shlomo (Cedars-Sinai) for the AtT-20 cell line. Finally, we thank Dr. David Vocadlo (Simon Fraser University) for sharing the OGT inhibitor 5S-GlcNAc.

## REFERENCES

1. Ezzat S, Asa SL, Couldwell WT, Barr CE, Dodge WE, Vance ML, McCutcheon IE. The prevalence of pituitary adenomas: A systematic review. Cancer 2004;101(3):613–619.

2. Dolecek TA, Propp JM, Stroup NE, Kruchko C. CBTRUS statistical report: Primary brain and central nervous system tumors diagnosed in the United States in 2005-2009. Neuro-Oncol. 2012;14(SUPPL.5). doi:10.1093/neuonc/nos218.

3. Gittleman H, Ostrom QT, Farah PD, Ondracek A, Chen Y, Wolinsky Y, Kruchko C, Singer J, Kshettry VR, Laws ER, Sloan AE, Selman WR, Barnholtz-Sloan JS. Descriptive epidemiology of pituitary tumors in the United States, 2004–2009: Clinical article. J. Neurosurg. 2014;121(3):527–535.

4. Fernandez A, Karavitaki N, Wass JAH. Prevalence of pituitary adenomas: a community-based, cross-sectional study in Banbury (Oxfordshire, UK). Clin. Endocrinol. (Oxf.) 2010;72(3):377–382.

5. Matta MP, Couture E, Cazals L, Vezzosi D, Bennet A, Caron P. Impaired quality of life of patients with acromegaly: Control of GH/IGF-I excess improves psychological subscale appearance. Eur. J. Endocrinol. 2008;158(3):305–310.

6. van der Klaauw AA, Kars M, Biermasz NR, Roelfsema F, Dekkers OM, Corssmit EP, Van Aken MO Havekes B, Pereira AM, Pijl H, Smit JW, Romijn JA. Disease-specific impairments in quality of life during long-term follow-up of patients with different pituitary adenomas. Clin. Endocrinol. (Oxf.) 2008;69(5):775–784.

7. Dekkers OM, Biermasz NR, Pereira AM, Roelfsema F, Van Aken MO, Voormolen JHC, Romijn JA. Mortality in patients treated for Cushing’s disease is increased, compared with patients treated for nonfunctioning pituitary macroadenoma. J. Clin. Endocrinol. Metab. 2007;92(3):976–981.

8. Asa SL, Ezzat S. The pathogenesis of pituitary tumours. Nat. Rev. Cancer 2002;2(11):836–849.

9. Rosén T, Bengtsson B-Å. Premature mortality due to cardiovascular disease in hypopituitarism. The Lancet 1990;336(8710):285–288.

10. Mohr G, Hardy J, Comtois R, Beauregard H. Surgical Management of Giant Pituitary Adenomas. Can. J. Neurol. Sci. J. Can. Sci. Neurol. 1990;17(1):62–66.

11. Selman WR, Laws ER, Scheithauer BW, Carpenter SM. The occurrence of dural invasion in pituitary adenomas. J. Neurosurg. 1986;64(3):402–407.

12. Meij BP, Lopes MBS, Ellegala DB, Alden TD, Laws ER. The long-term significance of microscopic dural invasion in 354 patients with pituitary adenomas treated with transsphenoidal surgery. J. Neurosurg. 2002;96(2):195–208.

13. Strasburger CJ, Karavitaki N, Störmann S, Trainer PJ, Kreitschmann-Andermahr I, Droste M, Korbonits M, Feldmann B, Zopf K, Sanderson VF, Schwicker D, Gelbaum D, Haviv A, Bidlingmaier M, Biermasz NR. Patient-reported outcomes of parenteral somatostatin analogue injections in 195 patients with acromegaly. Eur. J. Endocrinol. 2016;174(3):355–362.

14. Yamamoto M, Nakao T, Ogawa W, Fukuoka H. Aggressive Cushing’s Disease: Molecular Pathology and Its Therapeutic Approach. Front. Endocrinol. 2021;12:650791.

15. Gillam MP, Molitch ME, Lombardi G, Colao A. Advances in the treatment of prolactinomas. Endocr. Rev. 2006;27(5):485–534.

16. Hwang YC, Chung JH, Min YK, Lee MS, Lee MK, Kim KW. Comparisons between macroadenomas and microadenomas in cushing’s disease: Characteristics of hormone secretion and clinical outcomes. J. Korean Med. Sci. 2009;24(1):46–51.

17. Vandeva S, Vasilev V, Vroonen L, Naves L, Jaffrain-Rea M-L, Daly AF, Zacharieva S, Beckers A. Familial pituitary adenomas. Ann. Endocrinol. 2010;71(6):479–485.

18. Newey PJ, Nesbit MA, Rimmer AJ, Head RA, Gorvin CM, Attar M, Gregory L, Wass JAH, Buck D, Karavitaki N, Grossman AB, McVean G. Ansorge O, Thakker R V. Whole-exome sequencing studies of nonfunctioning pituitary adenomas. J. Clin. Endocrinol. Metab. 2013;98(4):796–800.

19. Németh K, Darvasi O, Likó I, Szücs N, Czirják S, Reiniger L, Szabó B, Kurucz PA, Krokker L, Igaz P, Patócs A, Butz H. Next-generation sequencing identifies novel mitochondrial variants in pituitary adenomas. J. Endocrinol. Invest. 2019;42(8):931–940.

20. Hauser BM, Lau A, Gupta S, Bi WL, Dunn IF. The epigenomics of pituitary adenoma. Front. Endocrinol. 2019;10(MAY):1–9.

21. Wulff-Fuentes E, Berendt RR, Massman L, Danner L, Malard F, Vora J, Kahsay R, Olivier-Van Stichelen S. The human O-GlcNAcome database and meta-analysis. Sci. Data 2021:1–11.

22. Hardivillé S, Hart GW. Nutrient regulation of signaling, transcription, and cell physiology by O-GlcNAcylation. Cell Metab. 2014;20(2):208–213.

23. Bolanle IO, Riches-Suman K, Williamson R, Palmer TM. Emerging roles of protein O-GlcNAcylation in cardiovascular diseases: Insights and novel therapeutic targets. Pharmacol. Res. 2021;165:105467.

24. Parker MP, Peterson KR, Slawson C. O-GlcNAcylation and O-GlcNAc Cycling Regulate Gene Transcription: Emerging Roles in Cancer. Cancers 2021;13(7):1666.

25. Park J, Lai MKP, Arumugam TV, Jo D-G. O-GlcNAcylation as a Therapeutic Target for Alzheimer’s Disease. Neuromolecular Med. 2020;22(2):171–193.

26. Hanover JA, Chen W, Bond MR. O-GlcNAc in cancer: An Oncometabolism-fueled vicious cycle. J. Bioenerg. Biomembr. 2018;50(3):155–173.

27. Fardini Y, Dehennaut V, Lefebvre T, Issad T. O-GlcNAcylation: A new cancer hallmark? Front. Endocrinol. 2013;4(AUG):1–14.

28. Makwana V, Ryan P, Patel B, Dukie SA, Rudrawar S. Essential role of O-GlcNAcylation in stabilization of oncogenic factors. Biochim. Biophys. Acta - Gen. Subj. 2019;1863(8):1302–1317.

29. Ortiz-Meoz RF, Merbl Y, Kirschner MW, Walker S. Microarray discovery of new OGT substrates: The medulloblastoma oncogene OTX2 is O-GlcNAcylated. J. Am. Chem. Soc. 2014;136(13):4845–4848.

30. Lu J, Montgomery BK, Chatain GP, Bugarini A, Zhang Q, Wang X, Edwards NA, Ray-Chaudhury A, Merrill MJ, Lonser RR, Chittiboina P. Corticotropin releasing hormone can selectively stimulate glucose uptake in corticotropinoma via glucose transporter 1. Mol. Cell. Endocrinol. 2018;470:105–114.

31. Staton JM, Thomson AM, Leedman PJ. Hormonal regulation of mRNA stability and RNA-protein interactions in the pituitary. J. Mol. Endocrinol. 2000;25(1):17–34.

32. Staton JM, Leedman PJ. Posttranscriptional regulation of thyrotropin beta-subunit messenger ribonucleic acid by thyroid hormone in murine thyrotrope tumor cells: a conserved mechanism across species. Endocrinology 1998;139(3):1093–1100.

33. Levy A, Lightman SL. Bromocriptine reduces rat thyrotropin beta-subunit mRNA stability. J. Endocrinol. Invest. 1990;13(1):49–53.

34. Das N, Kumar TR. Molecular regulation of follicle-stimulating hormone synthesis, secretion and action. J. Mol. Endocrinol. 2018;60(3):R131–R155.

35. Park D, Cheon M, Kim C, Kim K, Ryu K. Progesterone together with estradiol promotes luteinizing hormone beta-subunit mRNA stability in rat pituitary cells cultured in vitro. Eur. J. Endocrinol. 1996;134(2):236–242.

36. Chedrese PJ, Kay TW, Jameson JL. Gonadotropin-releasing hormone stimulates glycoprotein hormone alpha-subunit messenger ribonucleic acid (mRNA) levels in alpha T3 cells by increasing transcription and mRNA stability. Endocrinology 1994;134(6):2475–2481.

37. Ayoubi TA, Jenks BG, Roubos EW, Martens GJ. Transcriptional and posttranscriptional regulation of the proopiomelanocortin gene in the pars intermedia of the pituitary gland of Xenopus laevis. Endocrinology 1992;130(6):3560–3566.

38. Shan X, Vocadlo DJ, Krieger C. Reduced protein O-glycosylation in the nervous system of the mutant SOD1 transgenic mouse model of amyotrophic lateral sclerosis. Neurosci. Lett. 2012;516(2):296–301.

39. Massman L, Pereckas M, Zwagerman N, Olivier-Van Stichelen S. O-GlcNAcylation is essential for rapid Pomc expression and cell proliferation in corticotropic tumor cells (Supplementary Materials, Figures and Tables). 2021. doi:10.6084/m9.figshare15078270.v1.

40. Fabregat A, Sidiropoulos K, Viteri G, Marin-Garcia P, Ping P, Stein L, D’Eustachio P, Hermjakob H. Reactome diagram viewer: data structures and strategies to boost performance. Bioinforma. Oxf. Engl. 2018;34(7):1208–1214.

41. Ferrer CM, Sodi VL, Reginato MJ. O-GlcNAcylation in Cancer Biology: Linking Metabolism and Signaling. J. Mol. Biol. 2016;428(16):3282–3294.

42. Caldwell SA, Jackson SR, Shahriari KS, Lynch TP, Sethi G, Walker S, Vosseller K, Reginato MJ. Nutrient sensor O-GlcNAc transferase regulates breast cancer tumorigenesis through targeting of the oncogenic transcription factor FoxM1. Oncogene 2010;29(19):2831–2842.

43. Steenackers A, Olivier-Van Stichelen S, Baldini SF, Dehennaut V, Toillon R-A, Le Bourhis X, El Yazidi-Belkoura I, Lefebvre T. Silencing the Nucleocytoplasmic O-GlcNAc Transferase Reduces Proliferation, Adhesion, and Migration of Cancer and Fetal Human Colon Cell Lines. Front. Endocrinol. 2016;7:46.

44. Park H-J, Kim HJ, Lee JH, Lee JY, Cho BK, Kang JS, Kang H, Yang Y, Cho DH. Corticotropin-Releasing Hormone (CRH) Downregulates Interleukin-18 Expression in Human HaCaT Keratinocytes by Activation of p38 Mitogen-Activated Protein Kinase (MAPK) Pathway. J. Invest. Dermatol. 2005;124(4):751–755.

45. Refojo D, Echenique C, Müller MB, Reul JMHM, Deussing JM, Wurst W, Sillaber I, Paez-Pereda M, Holsboer F, Arzt E. Corticotropin-releasing hormone activates ERK1/2 MAPK in specific brain areas. Proc. Natl. Acad. Sci. U. S. A. 2005;102(17):6183–6188.

46. Malard F, Wulff-Fuentes E, Berendt RR, Didier G, Olivier-Van Stichelen S. Automatization and self-maintenance of the O-GlcNAcome catalog: a smart scientific database. Database 2021;2021(baab039). doi:10.1093/database/baab039.

47. Lewis BA, Burlingame AL, Myers SA. Human RNA Polymerase II Promoter Recruitment in Vitro Is Regulated by O-Linked N-Acetylglucosaminyltransferase (OGT). J. Biol. Chem. 2016;291(27):14056–14061.

48. Thomas MC, Chiang C-M. The general transcription machinery and general cofactors. Crit. Rev. Biochem. Mol. Biol. 2006;41(3):105–178.

49. Resto M, Kim B-H, Fernandez AG, Abraham BJ, Zhao K, Lewis BA. O-GlcNAcase Is an RNA Polymerase II Elongation Factor Coupled to Pausing Factors SPT5 and TIF1β. J. Biol. Chem. 2016;291(43):22703–22713.

50. Tan Z-W, Fei G, Paulo JA, Bellaousov S, Martin SES, Duveau DY, Thomas CJ, Gygi SP, Boutz PL, Walker S. O-GlcNAc regulates gene expression by controlling detained intron splicing. Nucleic Acids Res. 2020;48(10):5656–5669.

51. Colao A, di Somma C, Lombardi G, Pivonello R, di Sarno A. Dopamine receptor agonists for treating prolactinomas. Expert Opin. Investig. Drugs 2002;11(6):787–800.

52. Cooper O, Greenman Y. Dopamine Agonists for Pituitary Adenomas. Front. Endocrinol. 2018;9. doi:10.3389/fendo.2018.00469.

53. Zhao P, Hu W, Wang H, Yu S, Li C, Bai J, Gui S, Zhang Y. Identification of Differentially Expressed Genes in Pituitary Adenomas by Integrating Analysis of Microarray Data. Int. J. Endocrinol. 2015;2015:e164087.

54. Li M-D, Ruan H-B, Singh JP, Zhao L, Zhao T, Azarhoush S, Wu J, Evans RM, Yang X. O-GlcNAc Transferase Is Involved in Glucocorticoid Receptor-mediated Transrepression*,. J. Biol. Chem. 2012;287(16):12904–12912.

55. Ramirez DH, Aonbangkhen C, Wu H-Y, Naftaly JA, Tang S, O’Meara TR, Woo CM. Engineering a Proximity-Directed O-GlcNAc Transferase for Selective Protein O-GlcNAcylation in Cells. ACS Chem. Biol. 2020;15(4):1059–1066.

56. Olivier-Van Stichelen S, Guinez C, Mir A-M, Perez-Cervera Y, Liu C, Michalski J-C, Lefebvre T. The hexosamine biosynthetic pathway and O-GlcNAcylation drive the expression of β-catenin and cell proliferation. Am. J. Physiol. Endocrinol. Metab. 2012;302(4):E417–424.

57. Jin F, Yu C, Zhao D, Wu M, Yang Z. A correlation between altered O-GlcNAcylation, migration and with changes in E-cadherin levels in ovarian cancer cells. Exp. Cell Res. 2013;319(10):1482–1490.

58. Berthier A, Vinod M, Porez G, Steenackers A, Alexandre J, Yamakawa N, Gheeraert C, Ploton M, Maréchal X, Dubois-Chevalier J, Hovasse A, Schaeffer-Reiss C, Cianférani S, Rolando C, Bray F, Duez H, Eeckhoute J, Lefebvre T, Staels B, Lefebvre P. Combinatorial regulation of hepatic cytoplasmic signaling and nuclear transcriptional events by the OGT/REV-ERBα complex. Proc. Natl. Acad. Sci. U. S. A. 2018;115(47):E11033–E11042.

59. Yang WH, Kim JE, Nam HW, Ju JW, Kim HS, Kim YS, Cho JW. Modification of p53 with O-linked N-acetylglucosamine regulates p53 activity and stability. Nat. Cell Biol. 2006;8(10):1074–1083.

60. Moldovan IM, Şuşman S, Pîrlog R, Jianu EM, Leucuţa DC, Melincovici CS, Crişan D, Florian IŞ. Molecular markers in the diagnosis of invasive pituitary adenomas - an immunohistochemistry study. Romanian J. Morphol. Embryol. Rev. Roum. Morphol. Embryol. 2017;58(4):1357–1364.

61. Olivier-Van Stichelen S, Dehennaut V, Buzy A, Zachayus J-L, Guinez C, Mir A-M, El Yazidi-Belkoura I, Copin M-C, Boureme D, Loyaux D, Ferrara P, Lefebvre T. O-GlcNAcylation stabilizes β-catenin through direct competition with phosphorylation at threonine 41. FASEB J. Off. Publ. Fed. Am. Soc. Exp. Biol. 2014;28(8):3325–3338.

62. Queiroz RM, Chien J, Madan R, Slawson C, Dias W. O-GlcNAc Regulates p53 in Ovarian Cancer. FASEB J. 2016;30(S1):652.2–652.2.

63. Zhu Y, Hart GW. Targeting O-GlcNAcylation to develop novel therapeutics. Mol. Aspects Med. 2020:100885.

64. Grammatopoulos DK. Insights into mechanisms of corticotropin-releasing hormone receptor signal transduction. Br. J. Pharmacol. 2012;166(1):85–97.

65. Drouin J. 60 YEARS OF POMC: Transcriptional and epigenetic regulation of POMC gene expression. J. Mol. Endocrinol. 2016;56(4):T99–T112.

66. Yang X, Ongusaha PP, Miles PD, Havstad JC, Zhang F, So WV, Kudlow JE, Michell RH, Olefsky JM, Field SJ, Evans RM. Phosphoinositide signalling links O-GlcNAc transferase to insulin resistance. Nature 2008;451(7181):964–969.

67. Goldberg H, Whiteside C, Fantus IG. O-linked β-N-acetylglucosamine supports p38 MAPK activation by high glucose in glomerular mesangial cells. Am. J. Physiol. Endocrinol. Metab. 2011;301(4):E713–726.

68. Jiang M, Qiu Z, Zhang S, Fan X, Cai X, Xu B, Li X, Zhou J, Zhang X, Chu Y, Wang W, Liang J, Horvath T, Yang X, Wu K, Nie Y, Fan D. Elevated O-GlcNAcylation promotes gastric cancer cells proliferation by modulating cell cycle related proteins and ERK 1/2 signaling. Oncotarget 2016;7(38):61390–61402.

69. Zhang X, Ma L, Qi J, Shan H, Yu W, Gu Y. MAPK/ERK signaling pathway-induced hyper-O-GlcNAcylation enhances cancer malignancy. Mol. Cell. Biochem. 2015;410(1–2):101–110.

70. Zeidan Q, Hart GW. The intersections between O-GlcNAcylation and phosphorylation: implications for multiple signaling pathways. J. Cell Sci. 2010;123(Pt 1):13–22.

71. Yang X, Ongusaha PP, Miles PD, Havstad JC, Zhang F, So WV, Kudlow JE, Michell RH, Olefsky JM, Field SJ, Evans RM. Phosphoinositide signalling links O-GlcNAc transferase to insulin resistance. Nature 2008;451(7181):964–969.

72. Xie S, Jin N, Gu J, Shi J, Sun J, Chu D, Zhang L, Dai C-L, Gu J-H, Gong C-X, Iqbal K, Liu F. O-GlcNAcylation of protein kinase A catalytic subunits enhances its activity: a mechanism linked to learning and memory deficits in Alzheimer’s disease. Aging Cell 2016;15(3):455–464.

73. Erickson JR, Pereira L, Wang L, Han G, Ferguson A, Dao K, Copeland RJ, Despa F, Hart GW, Ripplinger CM, Bers DM. Diabetic hyperglycaemia activates CaMKII and arrhythmias by O-linked glycosylation. Nature 2013;502(7471):372–376.

74. Lee BE, Kim HY, Kim H-J, Jeong H, Kim B-G, Lee H-E, Lee J, Kim HB, Lee SE, Yang YR, Yi EC, Hanover JA, Myung K, Suh P-G, Kwon T, Kim J-I. O-GlcNAcylation regulates dopamine neuron function, survival and degeneration in Parkinson disease. Brain J. Neurol. 2020;143(12):3699–3716.

75. Dehennaut V, Slomianny M-C, Page A, Vercoutter-Edouart A-S, Jessus C, Michalski J-C, Vilain J-P, Bodart J-F, Lefebvre T. Identification of structural and functional O-linked N-acetylglucosamine-bearing proteins in Xenopus laevis oocyte. Mol. Cell. Proteomics MCP 2008;7(11):2229–2245.

76. Qin K, Zhu Y, Qin W, Gao J, Shao X, Wang Y, Zhou W, Wang C, Chen X. Quantitative Profiling of Protein O-GlcNAcylation Sites by an Isotope-Tagged Cleavable Linker. ACS Chem. Biol. 2018;13(8):1983–1989.

77. Huo B, Zhang W, Zhao X, Dong H, Yu Y, Wang J, Qian X, Qin W. A triarylphosphine-trimethylpiperidine reagent for the one-step derivatization and enrichment of protein post-translational modifications and identification by mass spectrometry. Chem. Commun. Camb. Engl. 2018;54(98):13790–13793.

78. Gao Y, Liu J, Bai Z, Sink S, Zhao C, Lorenzo FR, McClain DA. Iron down-regulates leptin by suppressing protein O-GlcNAc modification in adipocytes, resulting in decreased levels of O-glycosylated CREB. J. Biol. Chem. 2019;294(14):5487–5495.

79. Levine ZG, Fan C, Melicher MS, Orman M, Benjamin T, Walker S. O-GlcNAc Transferase Recognizes Protein Substrates Using an Asparagine Ladder in the Tetratricopeptide Repeat (TPR) Superhelix. J. Am. Chem. Soc. 2018;140(10):3510–3513.

80. Cassarino MF, Sesta A, Pagliardini L, Losa M, Lasio G, Cavagnini F, Pecori Giraldi F. Proopiomelanocortin, glucocorticoid, and CRH receptor expression in human ACTH-secreting pituitary adenomas. Endocrine 2017;55(3):853–860.

81. Zhang W, Liu T, Dong H, Bai H, Tian F, Shi Z, Chen M, Wang J, Qin W, Qian X. Synthesis of a Highly Azide-Reactive and Thermosensitive Biofunctional Reagent for Efficient Enrichment and Large-Scale Identification of O-GlcNAc Proteins by Mass Spectrometry. Anal. Chem. 2017;89(11):5810–5817.

82. Hahne H, Sobotzki N, Nyberg T, Helm D, Borodkin VS, van Aalten DMF, Agnew B, Kuster B. Proteome wide purification and identification of O-GlcNAc-modified proteins using click chemistry and mass spectrometry. J. Proteome Res. 2013;12(2):927–936.

83. Zhu Y, Willems LI, Salas D, Cecioni S, Wu WB, Foster LJ, Vocadlo DJ. Tandem Bioorthogonal Labeling Uncovers Endogenous Cotranslationally O-GlcNAc Modified Nascent Proteins. J. Am. Chem. Soc. 2020;142(37):15729–15739.

84. Woo CM, Lund PJ, Huang AC, Davis MM, Bertozzi CR, Pitteri SJ. Mapping and Quantification of Over 2000 O-linked Glycopeptides in Activated Human T Cells with Isotope-Targeted Glycoproteomics (Isotag). Mol. Cell. Proteomics MCP 2018;17(4):764–775.

85. Paul SJ, Ortolano GA, Haisenleder DJ, Stewart JM, Shupnik MA, Marshall JC. Gonadotropin subunit messenger RNA concentrations after blockade of gonadotropin-releasing hormone action: testosterone selectively increases follicle-stimulating hormone beta-subunit messenger RNA by posttranscriptional mechanisms. Mol. Endocrinol. Baltim. Md 1990;4(12):1943–1955.

86. Diamond DJ, Goodman HM. Regulation of growth hormone messenger RNA synthesis by dexamethasone and triiodothyronine: Transcriptional rate and mRNA stability changes in pituitary tumor cells. J. Mol. Biol. 1985;181(1):41–62.

87. Jones PM, Burrin JM. Ghatei MA, O’Halloran DJ, Legon S. Bloom SR. The Influence of Thyroid Hormone Status on the Hypothalamo-Hypophyseal Growth Hormone Axis*. Endocrinology 1990;126(3):1374–1379.

88. Imamura H, Wagih O, Niinae T, Sugiyama N, Beltrao P, Ishihama Y. Identifications of Putative PKA Substrates with Quantitative Phosphoproteomics and Primary-Sequence-Based Scoring. J. Proteome Res. 2017;16(4):1825–1830.

89. Lee S-H, Choi H-S, Kim H, Lee Y. ERK is a novel regulatory kinase for poly(A) polymerase. Nucleic Acids Res. 2008;36(3):803–813.

