## Supplementary Figures S1-S9 for "*O*-GlcNAcylation is essential for rapid *Pomc* expression and cell proliferation in corticotropic tumor cells"

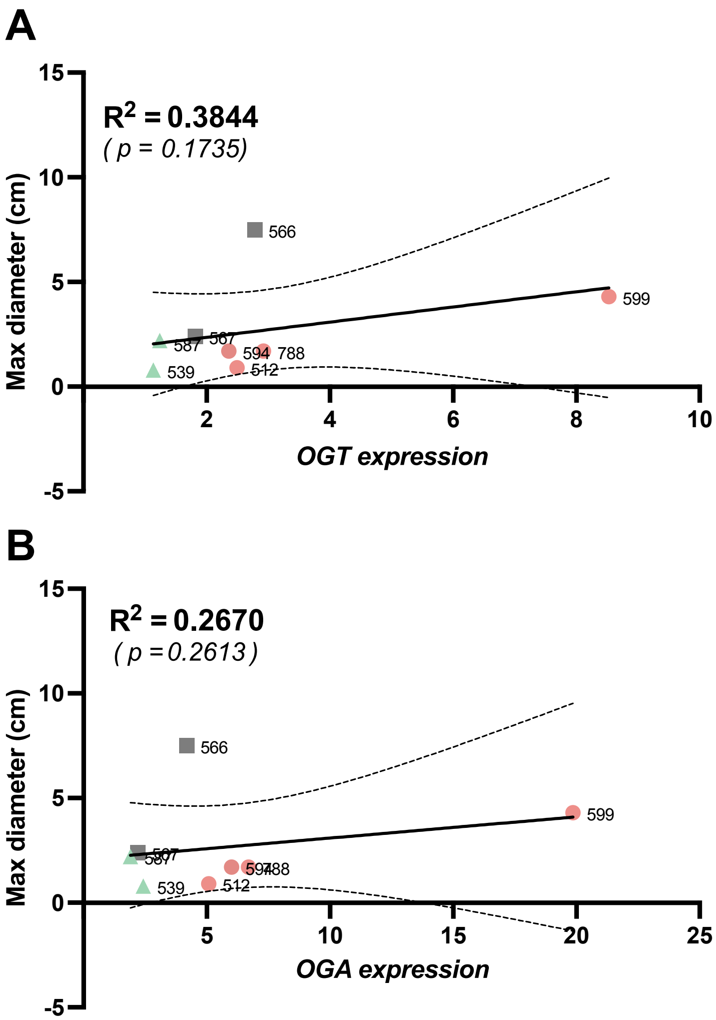


**Figure S1: Tumor size in pituitary adenomas does not correlate with *O*-GlcNAc enzyme expression. (A/B)** One-tailed Pearson correlations were computed between maximum diameter and (A) *OGT* and (B) *OGA* expression across pituitary adenoma subtypes ($R^2$ = 0.25, $p$ = *0.26* and $R^2$ = 0.38, $p$ = *0.17*, respectively). Simple linear regressions with 95% confidence intervals are represented.

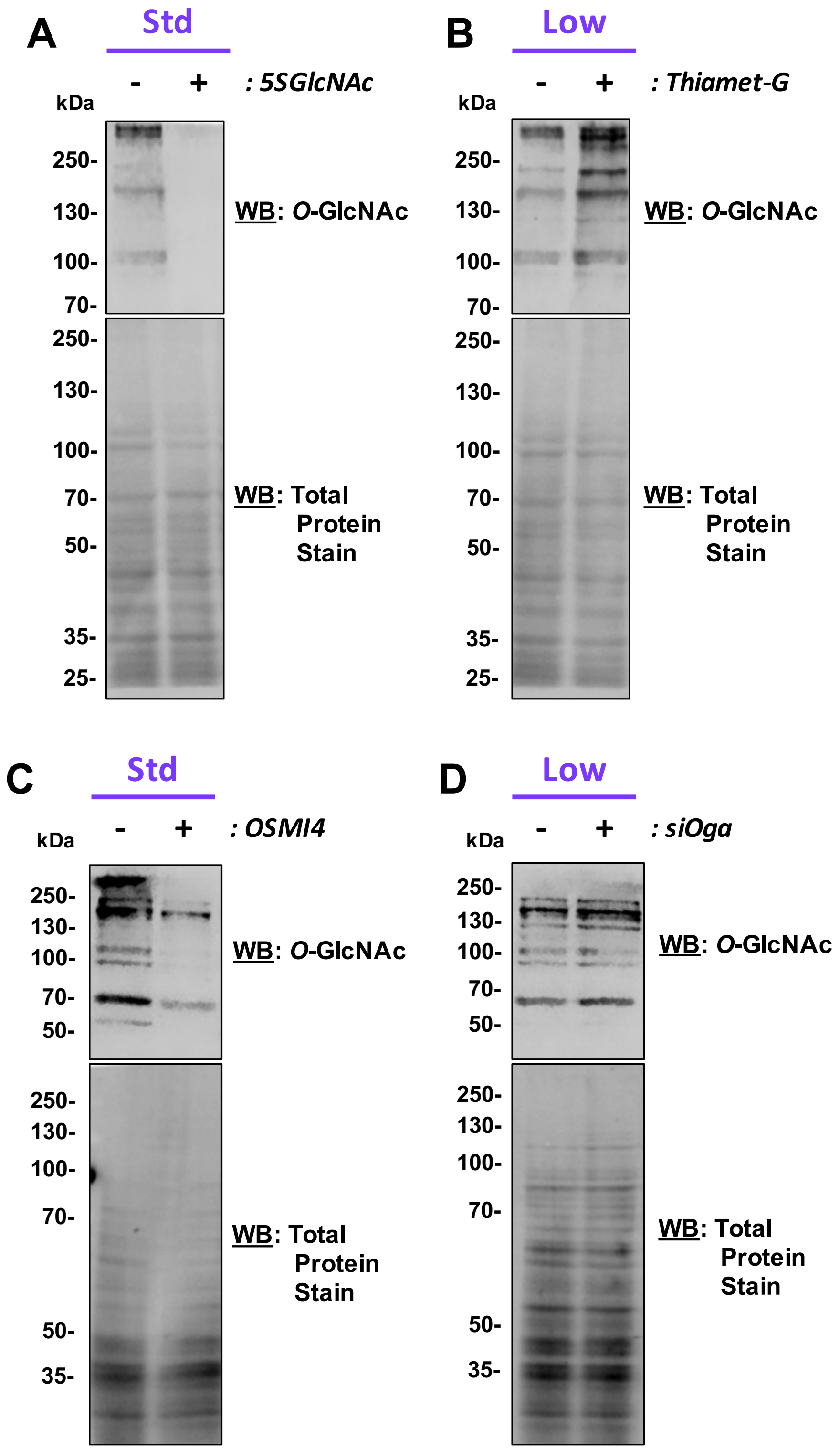


**Figure S2: O-GlcNAcylation levels are affected by glucose levels and inhibitor treatment in AtT-20 cells.** Cells were cultivated in **(A/C)** Standard (Std) or **(B/D)** low glucose media. Cells were treated overnight with 5S-GlcNAc **(A)**, Thiamet-G **(B)**, OSMI4 **(C)**, or transfected with siRNA against *Oga* **(D)**. Total *O*-GlcNAcylated protein levels were analyzed by western blot and normalized by Actin.

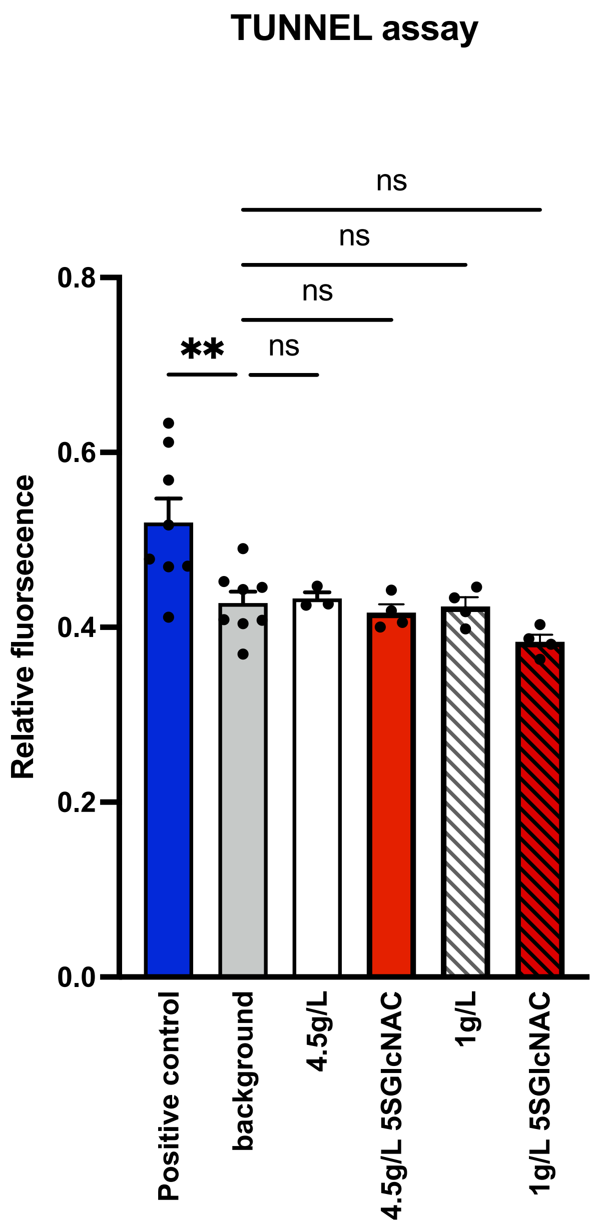


**<u>Figure S3:</u> Apoptotic rate of AtT-20 cells is not affected by 5S-GlcNAc treatment.** Apoptotic rate proliferation was measured by Tunel assay in triplicates for each condition. 5S-GlcNAc was added 24h prior to the assay into culture media, either standard (4.5g/L) or low (1g/L) glucose. Positive control cells were treated with DNAse.

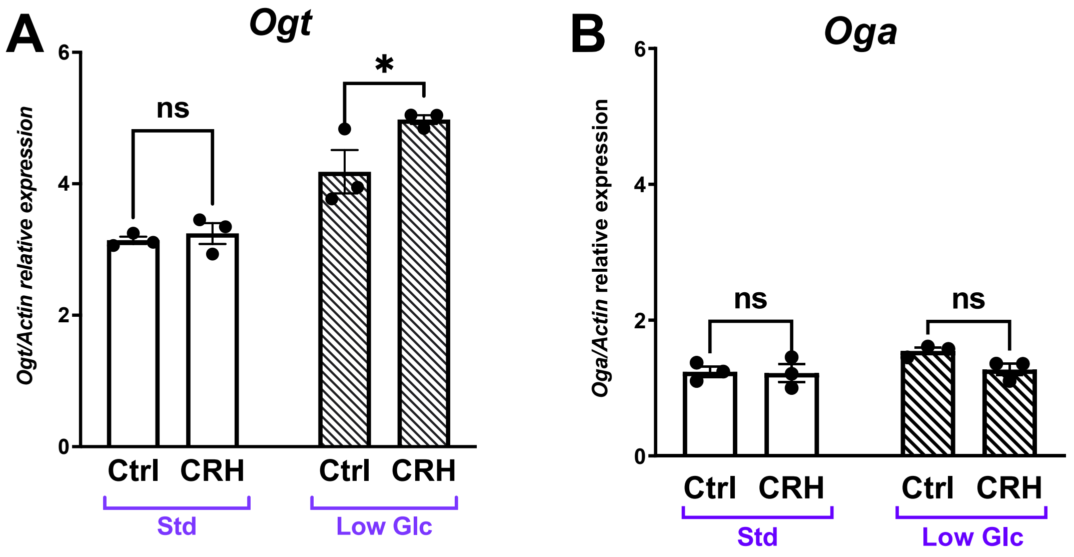


<u>Figure S4:</u> ***O*-GlcNAc enzymes are mildly altered by CRH stimulation (A/B)** qPCR for **(A)** expressions of *Ogt* and **(B)** *Oga* was analyzed by qPCR after CRH treatment (4h). Significance was measured by Two-way ANOVA with Fisher's LSD; *ns* $\geq 0.05$, * $p < 0.05$.

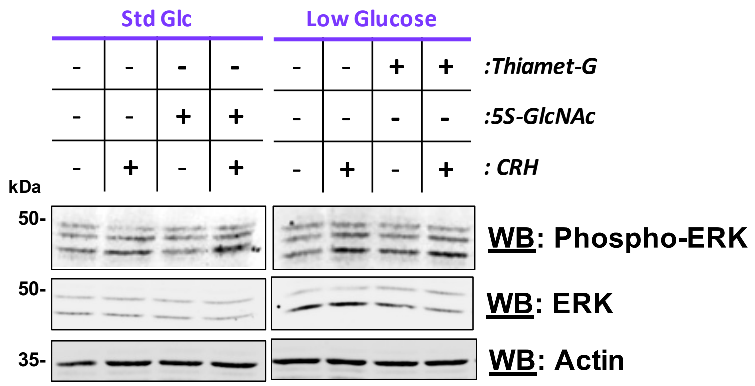


**Figure S5: Erk is stimulated at 30 min post-CRH treatment independently of *O*-GlcNAcylation status.** AtT-20 cells were treated with 5S-GlcNAc or Thiamet-G. Levels of phosphorylated Erk and total Erk were measured by western blot after CRH treatment (30min).

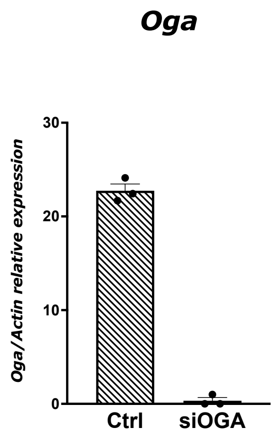


**<u>Figure S6</u>: *Oga* expression is reduced by transfection with *Oga* siRNA.** *Oga* levels measured by qPCR after 24h transfection with siRNA against *Oga* or negative control.

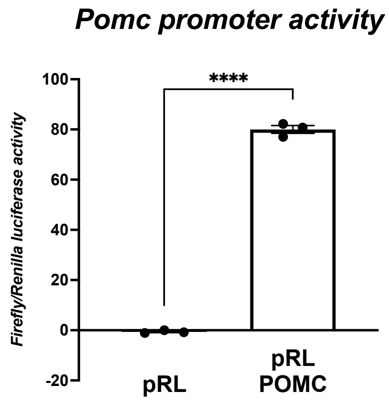


**<u>Figure S7:</u> Dual-luciferase assay is specific for the measurement of *Pomc* promoter activity.** AtT-20 cells were transfected with one or both of the plasmids (pRL-*Renilla* and *Pomc*-pGL3) used for the dual-luciferase reporter assay. Luciferase expression was measured using dual-luciferase assay protocol.

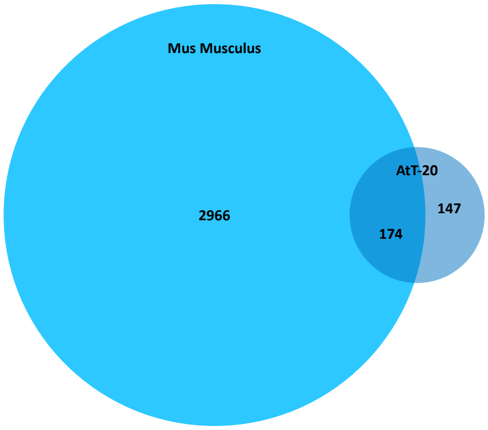


<u>**Figure S8**</u>**: Venn diagram of the *O*-GlcNAcomic analysis overlapping with the existing mouse *O*-GlcNAcome.** The existing list of *O*-GlcNAcylated protein in *Mus musculus* was extracted from the *O*-GlcNAc database.

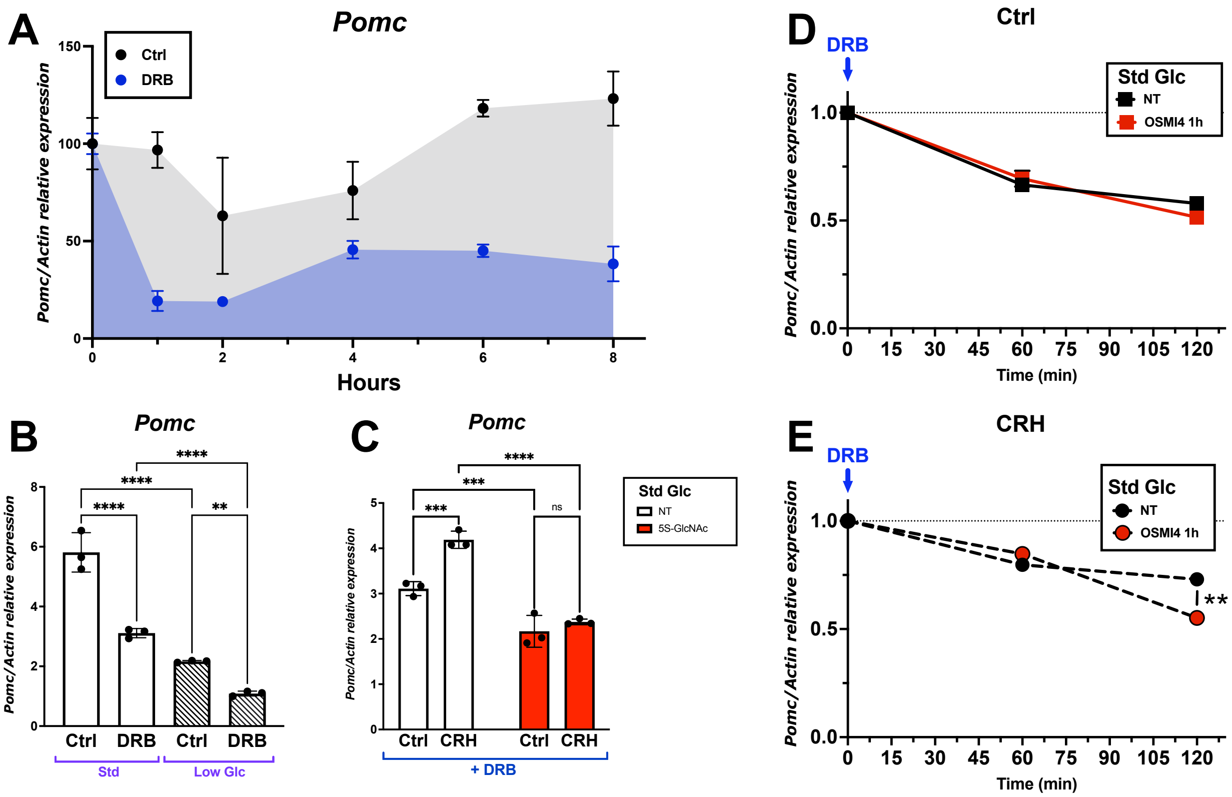


<u>**Figure S9:**</u> ***Pomc* transcripts are impacted by *O*-GlcNAcylation. (A)** Time-course of *Pomc* mRNA level was measured by qPCR following transcription inhibition by 5,6-dichloro-1-beta-D-ribofuranosysl-benzimidazole (DRB) in standard glucose condition. **(B)** *Pomc* mRNA levels in standard (Std) and low glucose conditions after one-hour DRB treatment. **(C)** *Pomc* levels at 1h post-DRB treatment on cells pre-treated with 5S-GlcNAc overnight in standard (Std) glucose media with or without CRH. **(D/E)** *Pomc* levels at 1h and 2h post-DRB treatment on cells pre-treated with OSMI4 for 1h in standard (Std) glucose media with or without CRH. Significance was measured by two-way ANOVA with Fisher's LSD; *ns ≥ 0.05, ** p < 0.01, *** p < 0.001, **** p < 0.0001*.
