## Supplementary Methods for "*O*-GlcNAcylation is essential for rapid *Pomc* expression and cell proliferation in corticotropic tumor cells"

**Supplementrary Methods.**

*Mass Spectrometry*

Each sample was analyzed on a Thermo Scientific Orbitrap Fusion Lumos MS via two technical replicate injections in each of two methods, one using HCD and product ion triggered EThcD (Table 1), the other HCD and EThcD for all $MS^2$ spectra (Table 2).

| **Table 1. Chromatography and MS instrument acquisition settings, 90 minute HCD/EThcD** | |
|---|---|
| ***Sample Volume*** | 10 µL |
| ***Stationary Phase*** | Thermo Acclaim PepMap $C_{18,}$ 75µm × 50cm |
| ***LC Solvent A*** | 100% $H_2O$, 0.1% formic acid |
| ***LC Solvent B*** | 80% acetonitrile, 0.1% formic acid |
| ***Column Temperature*** | 45°C |
| ***Gradient Ramp and Duration***<br>***Flow Rate*** | 1-3% B in 5 minutes<br>3-6% B in 5 minutes<br>6-10% B in 15 minutes<br>15-40% B in 10 minutes<br>40-99-99%B in 5 minutes<br>300 nL/min |
| ***Mass Spectrometer*** | Thermo Orbitrap Fusion Lumos |
| ***Spray Voltage*** | 2.2 kV |
| ***In-Source CID*** | 0.0 eV |
| ***RF Lens*** | 40% |
| ***Ion Transfer Tube Temp*** | 280 °C |
| ***$MS^1$ scan range*** | 300-2000 m/z |
| ***$MS^1$ resolution*** | 60,000 @ 200 *m/z* |
| ***$MS^1$ AGC Target*** | 2e5 |
| ***$MS^1$ Maximum IT*** | 100 ms |
| ***Isolation Window, HCD*** | 2 m/z |
| ***Charge state filter, HCD*** | 2-9, indetermined |
| ***Minimum Intensity Req. HCD*** | 2e4 |
| ***$MS^2$ AGC Target, HCD*** | 1e5 |
| ***$MS^2$ Maximum IT, HCD*** | 100 ms |
| ***Normalized Collision Energy, HCD*** | 15, 25, 35 Stepped |
| ***$MS^2$ Detection, HCD*** | Orbitrap, 15,000 @200 m/z, centroid |
| ***Charge state filter, EThcD*** | 2-9, |
| ***Minimum Intensity Req. EThcD*** | 2e4 |
| ***MS2 AGC Target, EThcD*** | 1e5 |
| ***MS2 Maximum IT, HCD*** | 100 ms |
| ***ETD Reaction time*** | Calibrated Charge-dependent, 10% SA energy |
| ***$MS^2$ Detection, EThcD*** | Orbitrap, 30,000 @ 200 m/z, profile, fixed 100-2000 m/z |
| ***Advanced Precursor Determination*** | off |
| ***Dynamic Exclusion*** | 20.0 s |
| ***MS2 acquisition*** | Data dependent, 3 s cycle time |

### Table 2. Chromatography and MS instrument acquisition settings, 180 minute HCD/Product Ion Triggered EThcD

| | |
|---|---|
| ***Sample Volume*** | 10 µL |
| ***Stationary Phase*** | Thermo Acclaim PepMap $C_{18}$, 75µm × 50cm |
| ***LC Solvent A*** | 100% $H_2O$, 0.1% formic acid |
| ***LC Solvent B*** | 80% acetonitrile, 0.1% formic acid |
| ***Column Temperature*** | 45°C |
| ***Gradient Ramp and Duration***<br>***Flow Rate*** | 1-1.5% B in 2 minutes<br>1.5-2% B in 3 minutes<br>2-3% B in 2 minutes<br>3-4% B in 3 minutes<br>4-5% B in 3 minutes<br>5-6% B in 2 minutes<br>6-9% B in 15 minutes<br>9-11% B in 15 minutes<br>11-13% B in 20 minutes<br>13-20% B in 15 minutes<br>20-22% B in 5 minutes<br>22-24% B in 5 minutes<br>24-26% B in 5 minutes<br>26-28% B in 5 minutes<br>28-30% B in 5 minutes<br>30-45% B in 10 minutes<br>45-99% B in 5 minutes<br>300 nL/min |
| ***Mass Spectrometer*** | Thermo Orbitrap Fusion Lumos |
| ***Spray Voltage*** | 2.35 kV |
| ***In-Source CID*** | 0.0 eV |
| ***RF Lens*** | 40% |
| ***Ion Transfer Tube Temp*** | 280 °C |
| ***$MS^1$ scan range*** | 200-2000 m/z |
| ***$MS^1$ resolution*** | 60,000 @ 200 *m/z* |
| ***$MS^1$ AGC Target*** | 2e5 |
| ***$MS^1$ Maximum IT*** | 100 ms |
| ***Isolation Window, HCD*** | 2 m/z |
| ***Charge state filter, HCD*** | 2-12, indetermined |
| ***Minimum Intensity Req. HCD*** | 3e4 |
| ***$MS^2$ AGC Target, HCD*** | 2.5e4 |
| ***$MS^2$ Maximum IT, HCD*** | 100 ms |
| ***Normalized Collision Energy, HCD*** | 15, 25, 35 Stepped |
| ***$MS^2$ Detection, HCD*** | Orbitrap, 30,000 @200 m/z, centroid |
| ***Targeted Mass Trigger Mass List, Minimum 2, ±10 ppm*** | 138.0549<br>168.0655<br>186.0761<br>204.0867 |
| ***MS2 AGC Target, EThcD*** | 1e5 |
| ***MS2 Maximum IT, HCD*** | 150 ms |
| ***ETD Reaction time*** | Calibrated Charge-dependent, 15% SA energy |
| ***$MS^2$ Detection, EThcD*** | Orbitrap, 30,000 @ 200 m/z, profile, fixed first mass 100 m/z |
| ***Advanced Precursor Determination*** | Off |
| ***Dynamic Exclusion*** | 20.0 s |
| ***MS2 acquisition*** | Data dependent, 5 s cycle time |

*Bioinformatic analysis*

MS data were analyzed using Proteome Discoverer 2.4 (Thermo) platform as outlined in the Table 3.

**Table 3. Mass spectrometry data processing parameters**

| | | | |
|---|---|---|---|
| ***Platform*** | ProteomeDiscoverer 2.4 | ***Static Modifications*** | Carbamidomethyl (C) |
| ***Search Algorithms*** | SequestHT | ***Dynamic Modifications*** | Oxidation (M), Acetylation (protein N-terminus) HexNAc (+203.079 on S, T, or Y) |
| ***Validation*** | Percolator, Protein FDR Validator | ***Target FDR (Strict) for PSMs:*** | 0.01 |
| ***Database*** | Swissprot human with isoforms, 2019-05-01, MaxQuant Contaminants | ***Target FDR (Relaxed) for PSMs:*** | 0.05 |
| ***Digest*** | Trypsin (semi)<br>2 Missed Cleavages Allowed | ***Target FDR (Strict) for Peptides:*** | 0.01 |
| ***Precursor mass tolerance*** | 10 ppm | ***Target FDR (Relaxed) for Peptides:*** | 0.05 |
| ***Fragment mass tolerance*** | 0.02 Da | | |
